# Resurrection of endogenous retroviruses during aging reinforces senescence

**DOI:** 10.1101/2021.02.22.432260

**Authors:** Xiaoqian Liu, Zunpeng Liu, Zeming Wu, Jie Ren, Yanling Fan, Liang Sun, Gang Cao, Yuyu Niu, Baohu Zhang, Qianzhao Ji, Xiaoyu Jiang, Cui Wang, Qiaoran Wang, Zhejun Ji, Lanzhu Li, Concepcion Rodriguez Esteban, Kaowen Yan, Wei Li, Yusheng Cai, Si Wang, Aihua Zheng, Yong E. Zhang, Shengjun Tan, Yingao Cai, Moshi Song, Falong Lu, Fuchou Tang, Weizhi Ji, Qi Zhou, Juan Carlos Izpisua Belmonte, Weiqi Zhang, Jing Qu, Guang-Hui Liu

## Abstract

Whether and how certain transposable elements with viral origins, such as endogenous retroviruses (ERVs) dormant in our genomes, can become awakened and contribute to the aging process are largely unknown. In human senescent cells, we found that HERVK (HML-2), the most recently integrated human ERVs, are unlocked to transcribe viral genes and produce retrovirus-like particles (RVLPs). These HERVK RVLPs constitute a transmissible message to elicit senescence phenotypes in young cells, which can be blocked by neutralizing antibodies. Activation of ERVs was also observed in organs of aged primates and mice, as well as in human tissues and serum from the elderly. Their repression alleviates cellular senescence and tissue degeneration and, to some extent, organismal aging. These findings indicate that the resurrection of ERVs is a hallmark and driving force of cellular senescence and tissue aging.

**In brief:** Liu and colleagues uncover the ways in which de-repression of human endogenous retrovirus triggers cellular senescence and tissue aging; the findings provide fresh insights into therapeutic strategies for alleviating aging.

**Highlights:**

- Derepression of the endogenous retrovirus contributes to programmed aging
- Upregulation of HERVK triggers the innate immune response and cellular senescence
- Extracellular HERVK retrovirus-like particles induce senescence in young cells
- Endogenous retrovirus serves as a potential target to alleviate agings

**Graphical abstract:** 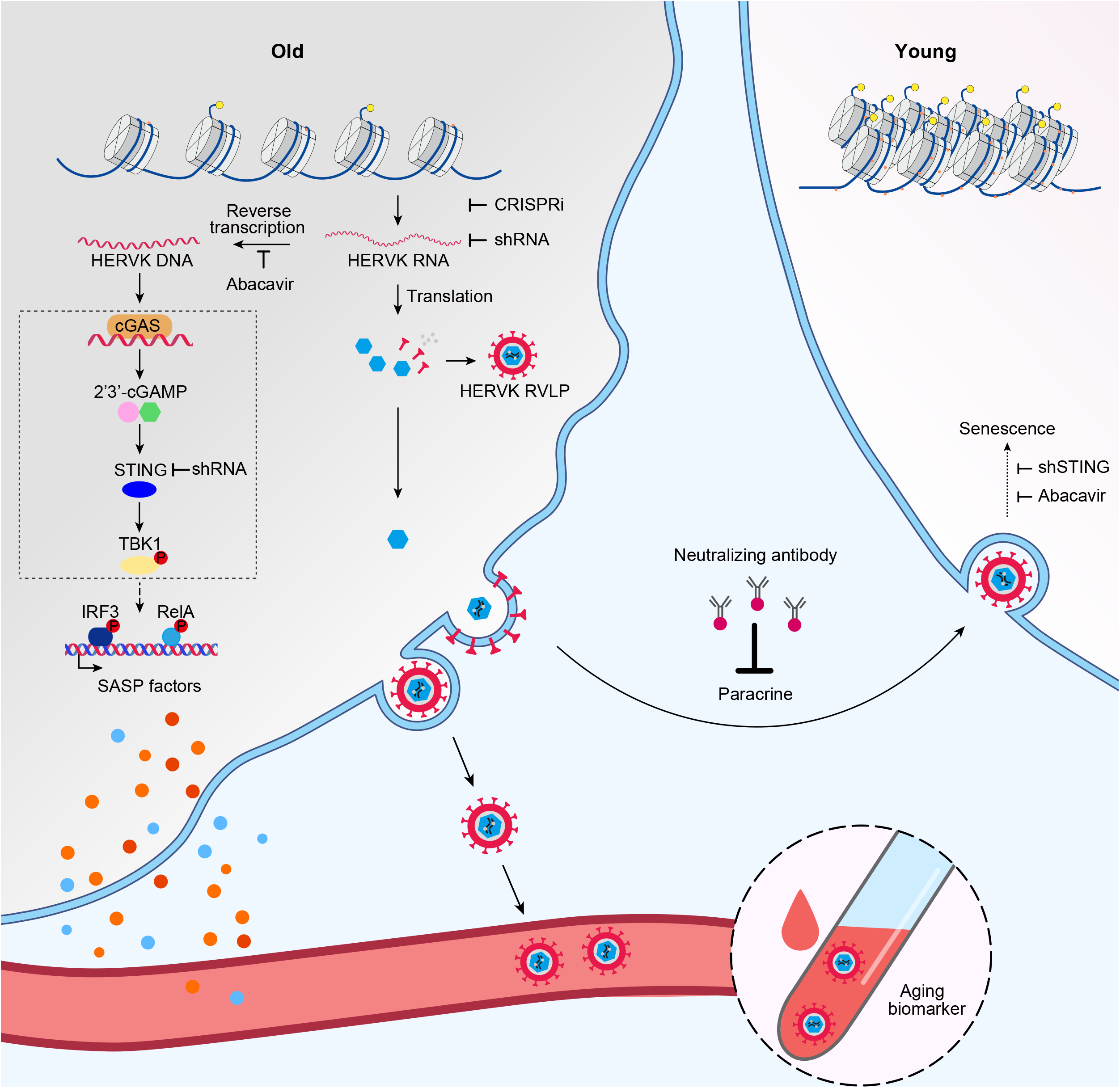

## INTRODUCTION

Aging is associated with a physiological decline and manifestations of chronic diseases, yet many of its underlying molecular changes and mechanisms remain poorly understood. With significant efforts over the past decades, several causal determinants of aging-related molecular changes have been identified, such as epigenetic alterations and stimulation of senescence-associated secretory phenotype (SASP) factors.^1–7^ Although the majority of these studies describe aging determinants originating primarily from protein-coding genes, the non-coding part of the genome has started to garner attention as well. For example, silent long-interspersed element-1 (LINE1) retrotransposons, belonging to non-long terminal repeat (non-LTR) retrotransposons, can be activated during senescence, triggering the innate immune response that is responsible for part of the senescence-associated phenotypes.^8–15^

A different class of retroelements, endogenous retroviruses (ERVs), belonging to LTR retrotransposons, are a relic of ancient retroviral infection, fixed in the genome during evolution, comprising about 8% of the human genome.^16–19^ As a result of evolutionary pressure, most human ERVs (HERVs) accumulate mutations and deletions that prevent their replication and transposition function.^20,21^ However, some evolutionarily young subfamilies of HERV proviruses, such as the recently integrated HERVK human mouse mammary tumor virus like-2 (HML-2) subgroup, maintains open reading frames encoding proteins required for viral particle formation, including Gag, Pol, Env, and Pro.^18,22^ Except at specific stages of embryogenesis when DNA is hypomethylated and under certain pathological conditions such as cancer,^23–27^ HERVs are transcriptionally silenced by host surveillance mechanisms such as epigenetic regulation in post-embryonic developmental stages.^28,29^ Notably, whether endogenous retroviruses can escape host surveillance during aging and, if so, what effects they may exert on cellular and organismal aging are still poorly investigated.

In this study, using cross-species models and multiple techniques, we revealed an uncharacterized role of endogenous retrovirus resurrection as a biomarker and driver for aging. Specifically, we identified endogenous retrovirus expression associated with cellular and tissue aging, and that the accumulation of HERVK retrovirus-like particles (RVLPs) mediates the aging-promoting effects in recipient cells. More importantly, we can inhibit endogenous retrovirus-mediated pro-senescence effects to alleviate cellular senescence and tissue degeneration *in vivo*, suggesting possibilities for developing therapeutic strategies to treat aging-related disorders.

## RESULTS

### Upregulated HERVK expression in senescent hMPCs is associated with epigenetic derepression

Cellular senescence is considered as a major contributing factor to aging and a hallmark of human progeroid diseases, i.e., Hutchinson–Gilford progeria syndrome (HGPS) and Werner syndrome (WS). ^2,30–36^ We previously demonstrated that HGPS human mesenchymal progenitor cells (hMPCs) *(LMNA*^G608G/+^ hMPCs) or WS hMPCs *(WRN^-/-^* hMPCs) recapitulated premature aging phenotypes (Figure 1A), characterized by increased senescence-associated β-galactosidase (SA-β-gal)-positive cells (Figure S1A). Reduced cellular proliferation rates were also observed in HGPS and WS hMPCs, as well as in replicatively senescent (RS) wild-type (WT) hMPCs, as evidenced by fewer Ki67-positive cells and decreased clonal expansion ability (Figures S1B–S1F).^31–35,37^

**Figure 1.**
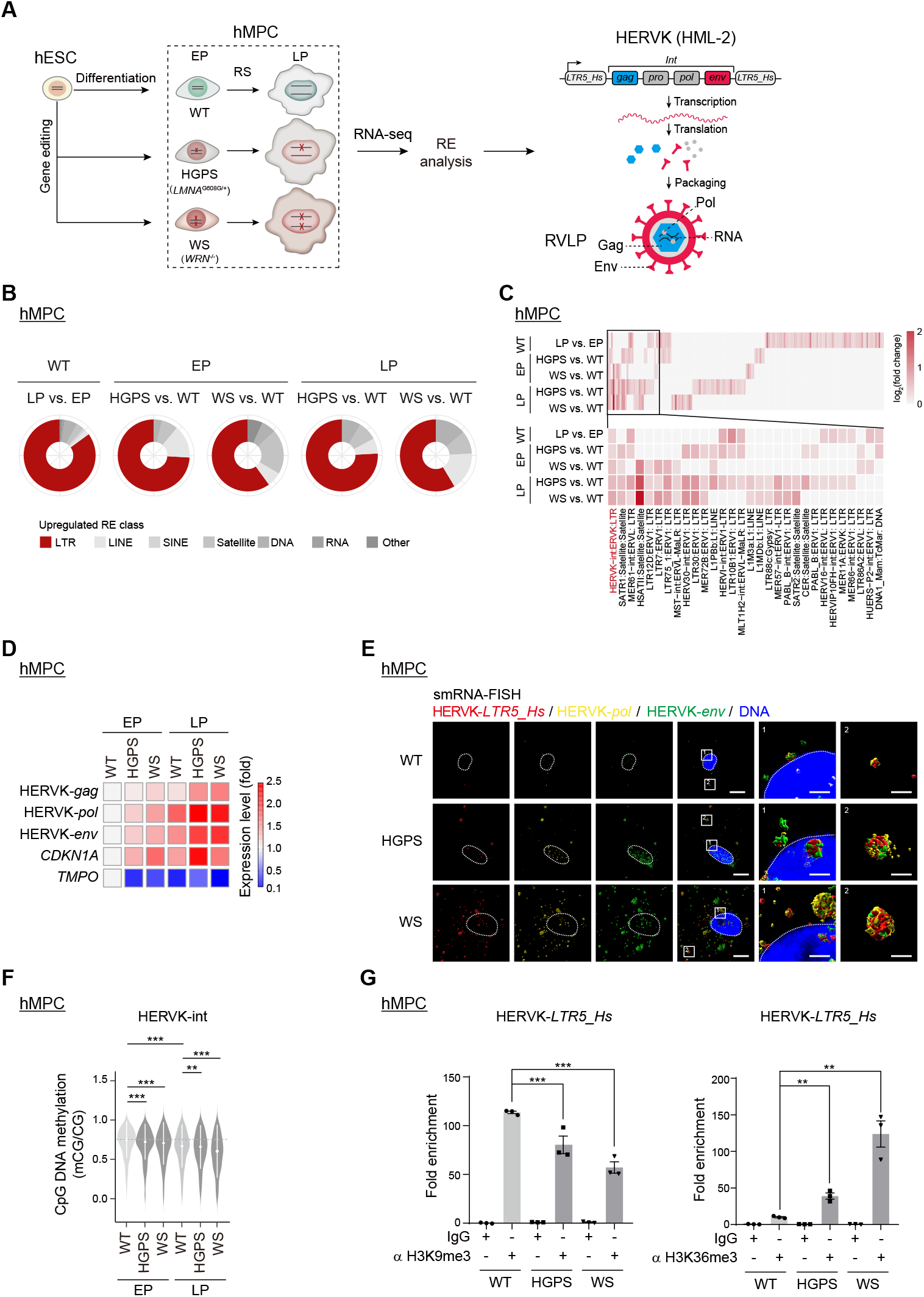
Epigenetic derepression of HERVK is observed in senescent hMPCs. (A) Schematic diagram of human aging stem cell models and the HERVK proviral genome structure. (B-C) Ring plots showing the percentages of upregulated repetitive elements in each class (B) and heatmap showing the relative expression levels for upregulated RepeatMasker-annotated repetitive elements (C) in RS and prematurely senescent hMPCs at early passage (EP) and late passage (LP). (D) Heatmap showing the levels of HERVK transcripts and senescence marker genes in WT, HGPS, and WS hMPCs at EP and LP as detected by qRT-PCR. (E) Representative Z-stack 3D reconstruction images of smRNA-FISH in WT, HGPS and WS hMPCs with probes targeting *HERVK-LTR5_Hs, -pol* and *-env* with different fluorophores. (F) Violin plot showing the CpG DNA methylation levels for HERVK-int in RS and prematurely senescent hMPCs. (G) ChIP-qPCR analysis of H3K9me3 and H3K36me3 enrichment in HERVK-*LTR5_Hs* regions in WT, HGPS, and WS hMPCs. Scale bars, 10 μm and 100 nm (zoomed-in image) in (E). See also Figure S1 and Table S1.

Here, we leveraged these premature aging models to test whether activation of the endogenous retroviruses is associated with human cellular senescence. We found increased expression of several transposable elements in senescent hMPCs, such as LTRs (Figure 1B and S1G; Table S1). Specifically, we found that retroelement HERVK internal coding sequences (HERVK-int) were upregulated in both replicatively and prematurely senescent hMPCs (Figures 1C and S1H; Table S1). Using primers targeting different regions of HERVK transcripts, including *env, pol*, and *gag*, we confirmed by quantitative reverse transcriptase PCR (qRT-PCR) that HERVK retroelements were highly expressed during cellular senescence, with a similar increase of the senescence marker p21^Cip1^ *(CDKN1A)*, while lamina-associated protein LAP2 *(TMPO)* decreased during senescence as reported previously (Figure 1D).^38^ Likewise, RNA fluorescence in situ hybridization (RNA-FISH) analysis also showed increased HERVK RNA signals in HGPS and WS hMPCs (Figure S1I). When we performed single molecule RNA-FISH (smRNA-FISH) with different fluorescent probes targeting *LTR5_Hs* (the transcriptional regulatory region of HERVK), *env*, and *pol*, we detected co-staining signals in close proximity, implying that mRNA molecules harboring *LTR5_Hs, env*, and *pol* are present in senescent hMPCs (Figure 1E).

Consistent with the increased expression of HERVK transcription in senescent hMPCs, we observed reduced CpG DNA methylation levels in HERVK-int regions and those HERVK proviral loci (Figures 1F, S1J and S1K),^39,40^ alongside a decrease in the repressive histone mark (H3K9me3) and an increase in the transcriptionally active histone mark (H3K36me3) at HERVK-*LTR5_Hs* (Figures 1G and S1L). These data indicate that the epigenetic derepression of HERVK, likely contributing to HERVK transcription, is associated with cellular senescence.

### Accumulation of viral proteins and RVLPs of HERVK in various types of senescent human cells

Next, we asked whether elevated endogenous retrovirus expression would elicit the production of HERVK protein components and even the formation of RVLPs. Western blotting and immunofluorescence staining analyses showed increased HERVK-Env protein levels in both prematurely senescent and RS hMPCs (Figures 2A, 2B, and S2A). Moreover, by co-staining HERVK-Env with SPiDER-βGal or p21^Cip1^, we found that the elevation of HERVK-Env was more pronounced in the senescent cell population (Figures S2B and S2C). In line with the increased levels of HERVK mRNA and protein, we detected an accumulation of RVLPs in the cytoplasm of both prematurely senescent and RS hMPCs by transmission electron microscopy (TEM) analysis; in contrast, RVLPs were very rare in early-passage WT hMPCs that were phenotypically young (Figures 2C and S2D). Using immuno-TEM analysis with an anti-HERVK-Env antibody,^41^ we found that these RVLPs, with a diameter ranging from 80 to 120 nm, were labeled with HERVK-Env antibody in senescent cells (Figures 2D, S2E, and S2F). These findings demonstrate the increased production of both viral proteins and RVLPs of HERVK in senescent hMPCs.

**Figure 2.**
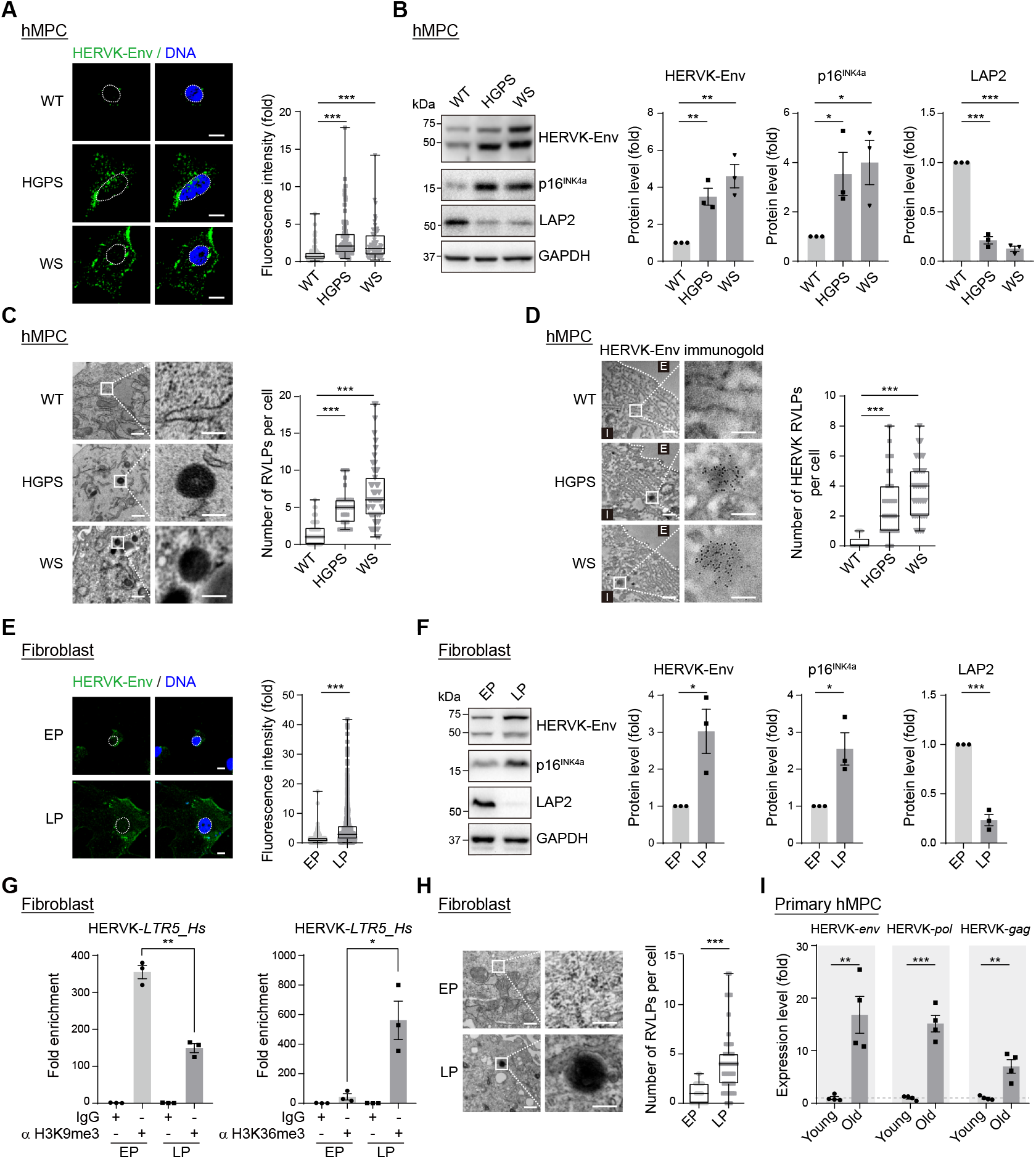
HERVK viral proteins and RVLPs are increased in senescent cells. (A) Immunofluorescence staining of HERVK-Env in WT, HGPS, and WS hMPCs. (B) Western blotting of HERVK-Env, p16^INK4a^, and LAP2 in WT, HGPS, and WS hMPCs. (C) TEM analysis of WT, HGPS and WS hMPCs. (D) TEM analysis after immunogold labeling with anti-HERVK-Env antibody in WT, HGPS, and WS hMPCs.: Extracellular; I:Intracellular. (E) Immunofluorescence staining of HERVK-Env in primary human fibroblasts at EP and LP. (F) Western blotting of HERVK-Env, p16^INK4a^, and LAP2 in primary human fibroblasts at EP and LP. (G) ChIP-qPCR analysis of H3K9me3 and H3K36me3 enrichment in HERVK-*LTR5_Hs* regions in primary human fibroblasts at EP and LP. (H) TEM analysis in primary human fibroblasts at EP and LP. (I) qRT-PCR analysis showing the relative expression of HERVK in primary hMPCs from young and old donors. Scale bars: 10 μm in (A) and (E); 200 nm and 100 nm (zoomed-in image) in (C), (D), and (H). See also Figure S2 and Table S2.

In order to verify the expression of HERVK in other senescent cell types, we isolated primary human fibroblasts and hMPCs. Similar to senescent hMPC models, we observed increased HERVK-Env protein levels in the replicatively senescent primary fibroblasts (Figures 2E, 2F and S2G–S2I). In addition, we found decreased occupancy of H3K9me3 and increased occupancy of H3K36me3 at HERVK-*LTR5_Hs* in human fibroblasts during replicative senescence (Figure 2G). We also observed an increased accumulation of RVLPs in these senescent primary fibroblasts (Figure 2H). Furthermore, we found prominent HERVK expression evidenced by a more than 5-fold increase in transcript levels in primary hMPCs derived from old individuals compared to their younger counterparts (Figures 2I and S2J; Table S2). Collectively, through multimodal experiments, we verified aberrant HERVK expression, as well as viral protein and RVLP accumulation in senescent cells.

### Increased expression of HERVK drives cellular senescence

To determine how activation of endogenous HERVK affects cellular senescence, we used a CRISPR-dCas9-mediated transcriptional activation (CRISPRa) system containing activation protein complexes (synergistic activation mediators, SAM) with sgRNAs targeting the HERVK-*LTR5_Hs* promoter regions in WT hMPCs (Figure S3A)^42^. We confirmed the activation of HERVK endogenous retrotransposon elements by qRT-PCR and western blotting (Figures 3A and S3B). We found that targeted HERVK activation induced hMPC senescence, as evidenced by the induction of classic senescent features (Figures 3B, 3C, S3B and S3C). To further assess whether suppression of endogenous HERVK inhibited cellular senescence, we used the CRISPR-dCas9-KRAB transcriptional inactivation (CRISPRi) system^43^ (Figures 3D, S3D, and S3E) to repress HERVK in prematurely senescent hMPCs and found that targeted HERVK repression alleviated hMPC senescence (Figures 3E, 3F, S3E, and S3F). In addition, the knockdown of HERVK using HERVK-interfering shRNA^44,45^ also antagonized premature senescence (Figures 3G, 3H, and S3G–S3I).

**Figure 3.**
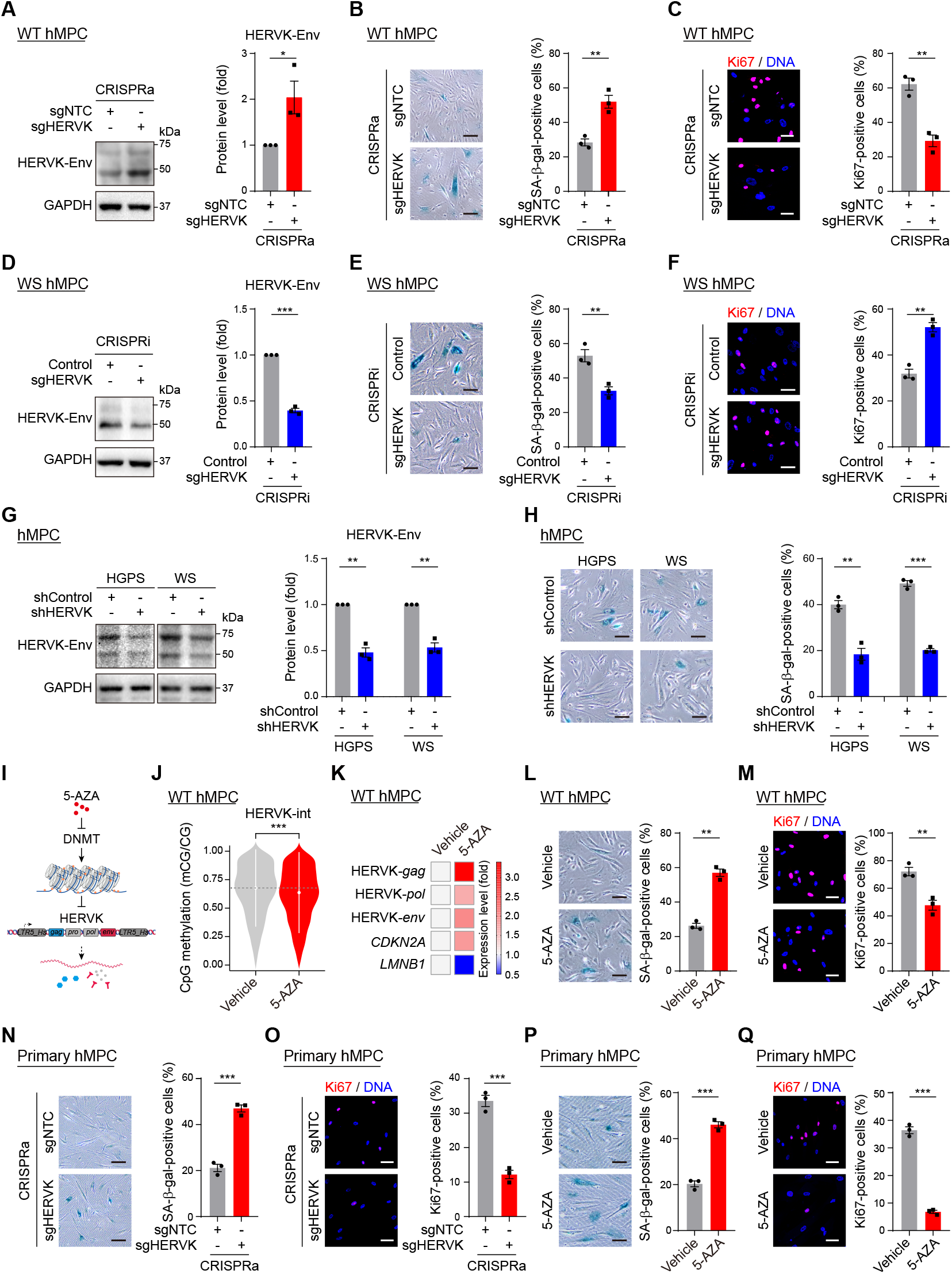
Increased HERVK drives senescence. (A-H) Western blotting of HERVK-Env, SA-β-gal and Ki67 staining of WT hMPCs transduced with lentiviruses expressing sgNTC or sgHERVK using a CRISPRa system (A-C), those expressing control or sgHERVK using a CRISPRi system (D-F), and those delivering shControl or shHERVK (G-H). (I) Schematic diagram showing the experimental design for activating HERVK with 5-AZA. (J) Violin plot showing the relative CpG DNA methylation levels for HERVK in WT hMPCs treated with vehicle or 5-AZA. (K) Heatmap showing the expression levels of *HERVK-gag/pol/env, CDKN2A*, and *LMNB1* in WT hMPCs treated with vehicle or 5-AZA by qRT-PCR analysis. (L-Q) SA-β-gal and Ki67 staining of WT hMPCs treated with vehicle or 5-AZA (L-M), of primary hMPCs from a young individual transduced with lentiviruses expressing sgNTC or sgHERVK using the CRISPRa system (N-O), and of vehicle- or 5-AZA-treated primary hMPCs from a young individual (P-Q). Scale bars, 20 μm (all panels). See also Figure S3.

Given the observed aging-associated decrease of DNA methylation at the HERVK loci in hMPCs (Figures 1F and S1K), we conducted an independent experiment in which we treated early-passage WT hMPCs with the DNA methyltransferase inhibitor (DNMTi) 5-azacytidine (5-AZA) to mimic aging-related global hypomethylation (Figure 3I).^46,47^ Whole-genome bisulfite sequencing (WGBS) analysis confirmed reduced CpG DNA methylation levels at HERVK elements after treatment (Figures 3J and S3J). Consistent with targeted HERVK activation by the CRISPRa system, 5-AZA treatment also led to upregulated HERVK RNA levels concomitant with premature cellular senescence phenotypes (Figures 3K–3M and S3K). In contrast, these senescence phenotypes were abrogated by HERVK knockdown (Figures S3L–S3N), suggesting that 5-AZA triggers cellular senescence at least partially via DNA demethylation-induced transcriptional activation of HERVK. Moreover, we also activated endogenous HERVK in primary hMPCs derived from a young individual via CRISPRa or 5-AZA treatment (Figures S3O and S3P) and found that transcriptional activation of HERVK accelerated cellular senescence (Figures 3N–3Q). Taken together, through disruption of the host epigenetic mechanism and targeted manipulation of HERVK transcriptional activity, we revealed that enhanced levels of endogenous HERVK are a driver of hMPC senescence.

### HERVK expression triggers the innate immune response

HERVK-encoded Pol protein possesses reverse transcription activity that can reverse-transcribe HERVK RNA into DNA,^16,17^ thereby generating additional HERVK DNA outside of the genome. Consistent with increased HERVK RNA (Figures 1D and S1I), we also observed increased HERVK DNA in the cytoplasm of senescent hMPCs by single-molecule DNA-FISH (Figures 4A and S4A). Therefore, we suspected that such excessive cytoplasmic DNA might be recognized by the DNA sensor cGMP-AMP synthase (cGAS) and trigger activation of the innate immune system (Figure 4B).^48–50^ Indeed, by immunoprecipitation analysis, we verified the marked enrichment of cGAS on cytoplasmic HERVK DNA in senescent hMPCs, which was not the case in early-passage young hMPCs (Figure 4C). Supporting cytosolic HERVK DNA triggering activation of the cGAS-Stimulator of interferon genes (STING) pathway, we detected increases in 2’3’-cGAMP content (Figure 4D), and phosphorylation of TANK-binding kinase 1 (TBK1), RelA and IFN regulatory factor 3 (IRF3) (Figure 4E). We also observed upregulation of inflammatory cytokines, including *IL1B* (IL1β) and *IL6*, both of which are classified as SASP factors^51,52^ (Figures 4F, 4G, and S4B; Table S3) in prematurely senescent hMPCs. Activation of the cGAS-STING-mediated innate immune response was also revealed in replicatively senescent hMPCs (Figures S4C–S4E; Table S3) and replicatively senescent fibroblasts (Figures S4F–S4H).

**Figure 4.**
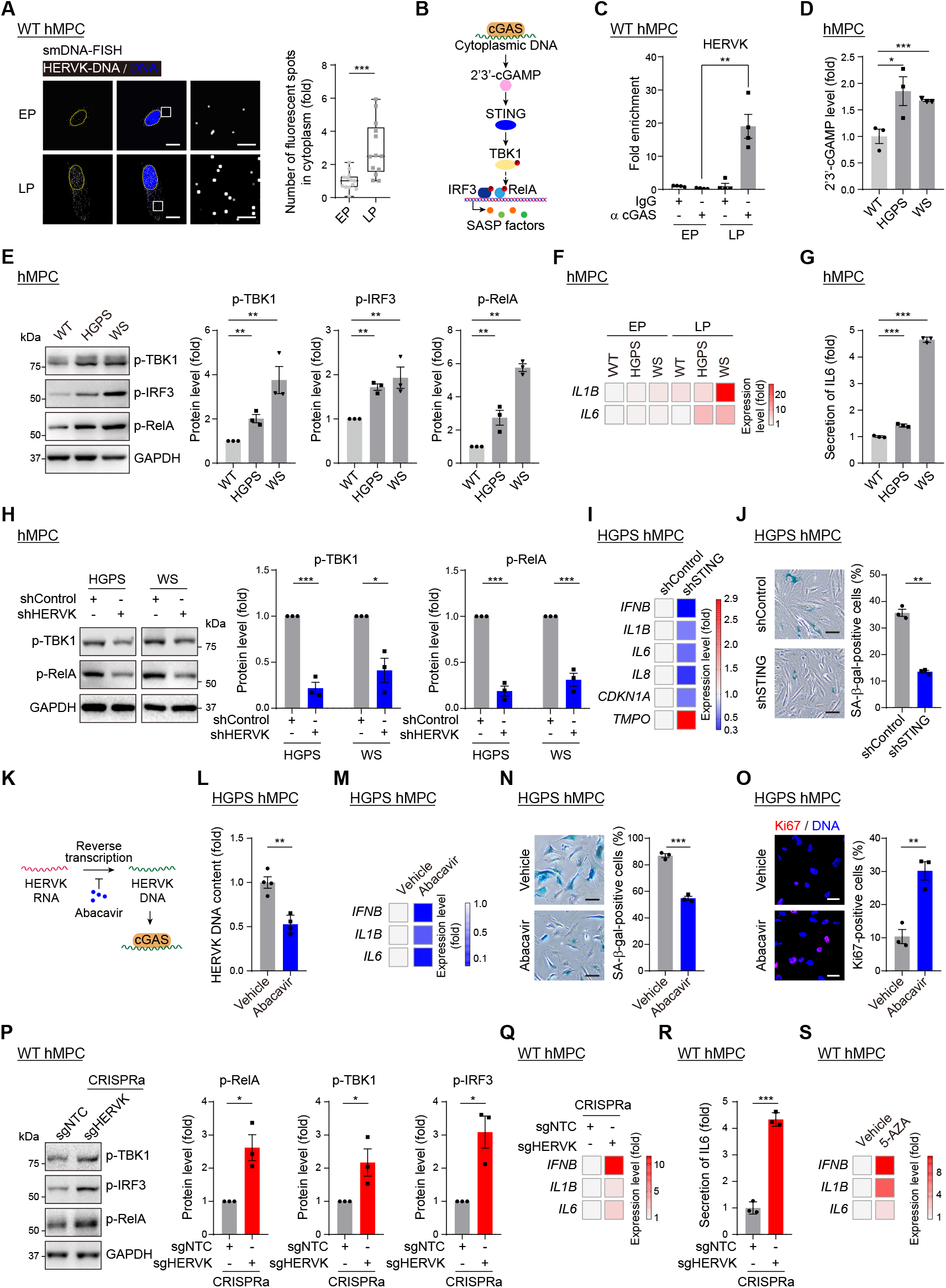
Increased HERVK activates the innate immunity pathway. (A) HERVK smDNA-FISH in RS WT hMPCs. (B) Schematic diagram showing the innate immune response through the cGAS-STING pathway. (C) Immunoprecipitation assay followed by qPCR analysis to assess cGAS enrichment on cytoplasmic HERVK DNA fragments in RS WT hMPCs. (D-G) ELISA analysis of 2’3’-cGAMP levels (D), western blotting of p-TBK1, p-IRF3, and p-RelA (E), qRT-PCR analysis the levels of SASP genes (F), and ELISA analysis of IL6 levels in the culture medium (G), in WT, HGPS, and WS hMPCs. (H) Western blotting of p-TBK1 and p-RelA in HGPS or WS hMPCs after transduction with lentiviruses delivering shControl or shHERVK. (I-J) qRT-PCR analysis of the levels of SASP and senescence marker genes (I), and SA-β-gal staining (J) of HGPS hMPCs after transduction with lentiviruses expressing shControl or shSTING. (K) Schematic diagram showing the experimental design for repressing HERVK with Abacavir. (L-O) qPCR analysis of the HERVK DNA contents (L) and qRT-PCR analysis of the expression of SASP genes (M), as well as SA-β-gal (N) and Ki67 (O) staining of HGPS hMPCs treated with vehicle or Abacavir. (P-R) Western blotting showing the protein levels of p-TBK1, p-IRF3, and p-RelA (P), as well as qRT-PCR analysis of the expression of SASP genes (Q), and ELISA analysis of IL6 levels in the culture medium (R), in WT hMPCs transduced with lentiviruses expressing sgNTC or sgHERVK using a CRISPRa system. (S) Heatmap showing qRT-PCR analysis of the levels of SASP genes in hMPCs treated with vehicle or 5-AZA. Scale bars: 10 μm and 200 nm (zoomed-in images) in (A); and 20 μm in (J), (N), and (O). See also Figure S4 and Tables S3 and S4.

Consistently, inhibition of HERVK via shRNA decreased the phosphorylation levels of TBK1 and RelA (Figure 4H). In addition, blocking one downstream effector of HERVK activation via knockdown of STING reduced both the inflammatory response and SASP expression (Figures 4I and S4I), thus alleviating cellular senescence in prematurely senescent hMPCs (Figure 4J). Moreover, we also blocked the downstream effect through Abacavir, a potent nucleoside reverse transcriptase inhibitor that can inhibit the activity of HERVK-encoded Pol (Figure 4K).^53^ Senescent hMPCs treated with Abacavir demonstrated diminished HERVK DNA content (Figure 4L), along with substantial alleviation of a panel of senescence-associated phenotypes (Figures 4M–4O, S4J, and S4K). In contrast to the loss-of-function experiments described above, activation of endogenous HERVK via the CRISPRa system or 5-AZA treatment led to an augmented innate immune response and upregulated expression of SASP cytokines in young WT hMPCs (Figures 4P–4S, S4L, and S4M; Table S4). These findings place the increased expression of HERVK as a contributing factor for cellular senescence, at least in part by triggering innate immune responses.

### Extracellular HERVK RVLPs induce cellular senescence

Previous studies indicated that tumor cell-derived HERVK RVLPs could be released into the culture medium and then taken up by other cells.^54^ Given that the presence of HERVK RVLPs in senescent cells was observed in this study (Figures 2C, 2D, and S2D–S2F), we asked whether HERVK RVLPs produced by senescent cells could be released extracellularly and convey senescence signals to nonsenescent cells (Figure 5A). To answer this question, we employed the sensitive droplet digital PCR (ddPCR) technology^55^ to detect HERVK RNA (the genetic material that is supposed to be packaged in RVLPs) in conditioned medium (CM) harvested from WT and prematurely senescent hMPCs. In CM from prematurely senescent hMPCs, we found that the HERVK RNA level was 5~12 times higher than that in CM from young WT hMPCs (Figure 5B). In addition, both enzyme-linked immunosorbent assay (ELISA) and western blotting demonstrated increased HERVK-Env protein levels in CM from prematurely senescent hMPCs (Figures 5C and S5A), as well as in CM from replicatively senescent hMPCs (Figures S5B and S5C). Furthermore, through TEM and immuno-TEM analyses, we detected RVLPs with diameters spanning from 80 to 120 nm, primarily on the outside of senescent hMPCs; some HERVK virial particles were detected budding from or adjacent to the cell surface (Figures 5D and S5D), whereas some particles were contained within coated vesicles in the extracellular environment (Figures S5D and S5E).

**Figure 5.**
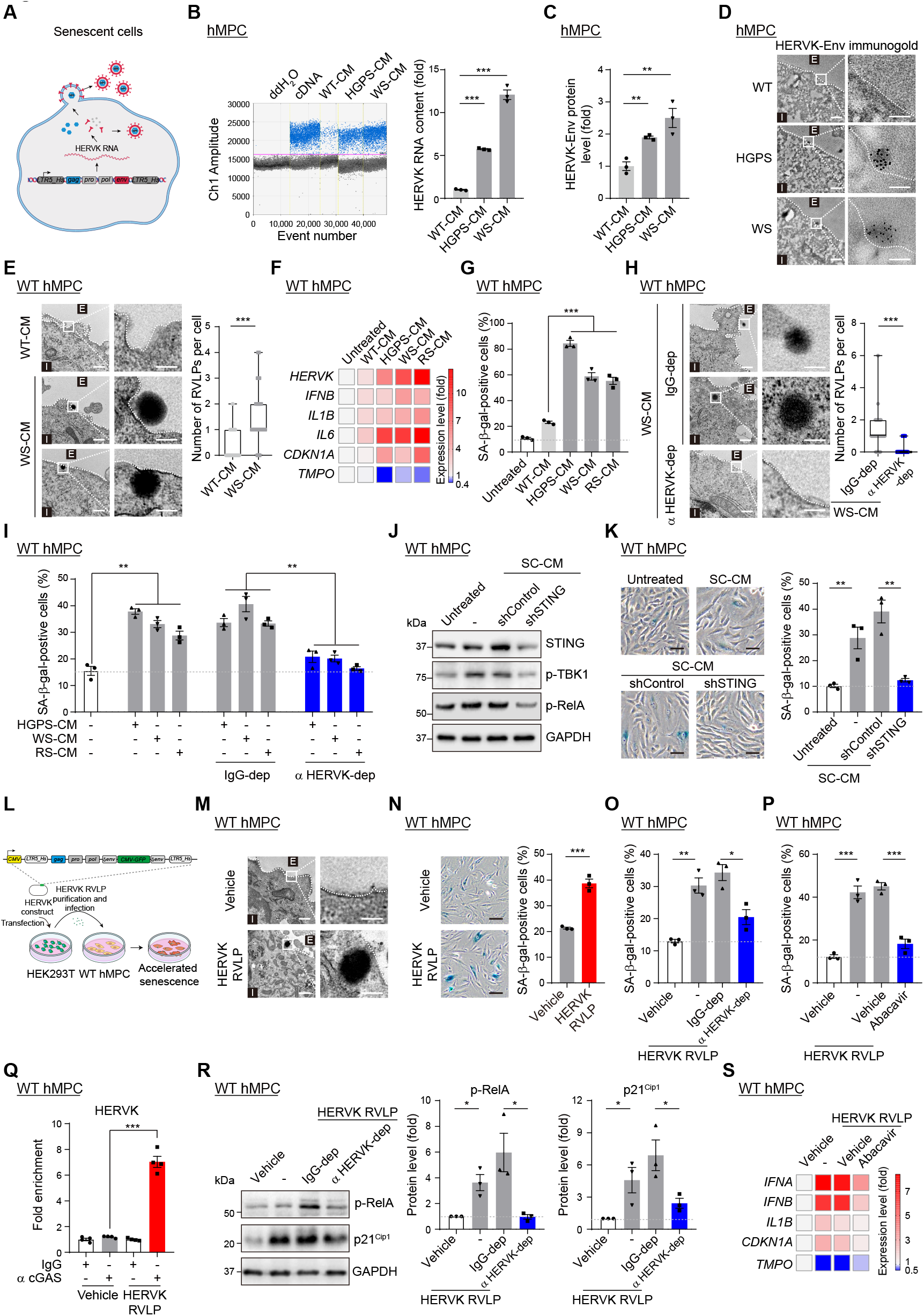
HERVK RVLPs released by senescent cells induce senescence in young cells. (A) Schematic diagram showing the proposed HERVK viral life cycle in senescent cells. (B-C) Digital droplet PCR (ddPCR) analysis of HERVK RNA levels (B) and ELISA analysis of HERVK-Env protein levels (C) in the CM of WT, HGPS, and WS hMPCs. (D) TEM analysis after immunogold labeling with anti-HERVK-Env antibody in WT, HGPS and WS hMPCs. (E) TEM analysis of young hMPCs treated with CM from young WT or senescent WS hMPCs. (F-G) qRT-PCR analysis of the levels of *HERVK*, senescence marker genes and inflammatory cytokines (F), as well as statistical analysis of SA-β-gal-positive cells (G) in young hMPCs treated with SC-CM. (H-I) TEM analysis (H) and statistical analysis of SA-β-gal-positive cells (I) in young hMPCs treated with SC-CM after immunodepletion with IgG or anti-HERVK-Env antibody. (J-K) Western blotting of STING, p-TBK1, and p-RelA (J), as well as SA-β-gal staining (K) in young hMPCs treated with SC-CM after STING knockdown. (L) Schematic diagram showing the experimental procedure for HERVK RVLP infection of WT hMPCs. (M-P) TEM images (M) and SA-β-gal staining (N) of WT hMPCs infected with HERVK RVLPs, as well as SA-β-gal staining of WT hMPCs infected with HERVK RVLPs after pretreatment with IgG or anti-HERVK-Env antibody (O), or in the presence of Abacavir (P). (Q) Immunoprecipitation assay followed by qPCR analysis to assess cGAS enrichment on HERVK DNA fragments in WT hMPCs infected with HERVK RVLPs. (R) Western blotting of p-RelA and p21^Cip1^ in WT hMPCs infected with HERVK RVLPs after pretreatment with IgG or anti-HERVK-Env antibdoy. (S) Heatmap showing the qRT-PCR analysis of the expression levels of inflammatory cytokines and senescence marker genes in WT hMPCs infected with HERVK RVLPs in the presence of Abacavir. Scale bars: 200 nm and 100 nm (zoomed-in images) in (D), (E), (H), and (M); and 20 μm in (K) and (N). See also Figure S5.

We next treated young hMPCs (WT) with CM collected from HGPS, WS, or RS hMPCs (referred to as senescent cell-conditional medium or SC-CM), using CM collected from young WT hMPCs as a control. With TEM analysis, we found that more extracellular RVLPs adhered to the cell surface of young hMPCs or entered the young cells in the SC-CM-treated groups (Figure 5E). After SC-CM treatment, we observed an increased HERVK abundance in young hMPCs (Figure 5F), implying that HERVK in SC-CM may be transmitted into target cells. Moreover, we found that the invaded HERVK elements were associated with an “aging-promoting” effect; that is, young hMPCs that were incubated with SC-CM also underwent accelerated cellular senescence (Figures 5F, 5G, and S5F).

To further investigate whether HERVK present in SC-CM is one of the major factors that causes senescence in young hMPCs, we used an anti-HERVK-Env antibody^56^ to immunodeplete HERVK. Western blotting confirmed the pulldown of both HERVK-Env and Gag proteins in SC-CM after incubation with the anti-HERVK-Env antibody, but not in SC-CM incubated with the IgG control (Figure S5G). Accordingly, the ELISA data showed depletion of HERVK-Env from SC-CM after incubation with the anti-HERVK-Env antibody (Figure S5H). As expected, after HERVK immunodepletion, we observed that fewer HERVK RVLPs adhered to the surface or were present in young hMPCs (Figure 5H). Furthermore, immunodepletion of HERVK resulted in reduction of HERVK abundance and alleviation of senescence phenotypes in young WT hMPCs compared to those treated with SC-CM without immunodepletion (Figures 5I, S5I, and S5J). Moreover, SC-CM activated the cGAS-STING pathway and induced the expression of SASP genes in young hMPCs (Figures 5F, 5J, and S5K). Importantly, knockdown of STING rescued the senescent phenotypes induced by SC-CM (Figures 5J, 5K, and S5K), indicating that SC-CM, similar to endogenous HERVK expression, drove cellular senescence at least partially by activating the innate immunity pathway. Collectively, these data suggest that retroviral HERVK elements generated in senescent cells can be released in a paracrine manner and trigger cellular senescence in non-senescent cells.

Next, in order to verify that the infection of extracellular HERVK RVLPs is a direct driver of senescence, we employed our constructed expression vector containing synthesized full-length HERVK with a GFP cassette fused within Env to produce RVLPs (Figures S5L and S5M).^57,58^ We then infected young hMPCs with purified HERVK RVLPs (Figures 5L and 5M), and found that both the RNA levels of HERVK and GFP fragments were upregulated in young hMPCs (Figure S5N). Young hMPCs infected with HERVK RVLPs, similar to those treated with SC-CM, showed typical premature senescence phenotypes (Figure 5N). And resembling SC-CM with immunodepletion, HERVK RVLP neutralization with the anti-HERVK-Env antibody abrogated HERVK RVLP-induced cellular senescence (Figures 5O, S5O, and S5P); such abrogation was also achieved by Abacavir treatment (Figures 5P, S5Q, and S5R). We further found an increased enrichment of cGAS on HERVK DNA in recipient hMPCs after infection with HERVK RVLPs (Figure 5Q). Additionally, we also detected activation of cGAS-STING-mediated innate immune responses in infected hMPCs (Figures 5R, 5S, and S5S). Of note, these phenotypes were abrogated by neutralization with an anti-HERVK-Env antibody or treatment with Abacavir (Figures 5R and 5S), demonstrating that the cGAS-STING pathway drives the acquisition of senescence caused by HERVK RVLPs. Taken together, our data support that extracellular HERVK RVLPs transmit aging information to young cells.

### Endogenous retrovirus can be used as a biomarker of aging

Next, we measured the levels of endogenous retroviral elements in aged primates and human individuals. ERVW is another endogenous retrovirus subfamily that persists in Old World monkeys due to infection of the ancestral germline when apes and Old World monkeys diverged about 25 million years ago.^59,60^ RT-qPCR results revealed that both ERVK and ERVW elements were increased in aged cynomolgus monkeys (Figure S6A), and with available antibodies,^61^ the ERVW-Env protein levels were confirmed to increase in lung, liver, and skin tissues in physiologically aged cynomolgus monkeys relative to younger counterparts (Figures 6A–6C). Consistently, we also found that the innate immune responses in aged monkey tissues were increased, as evidenced by the upregulation of p-RelA (Figure 6B) and SASP factors (Figures S6B–S6D). Furthermore, when comparing lung, liver, and skin tissues isolated from both WT and HGPS cynomolgus monkeys that recapitulated the premature aging phenotypes of HGPS patients,^62^ we found that the protein levels of ERVW-Env, along with p-RelA, were also increased in HGPS cynomolgus monkeys (Figures 6D–6F).

**Figure 6.**
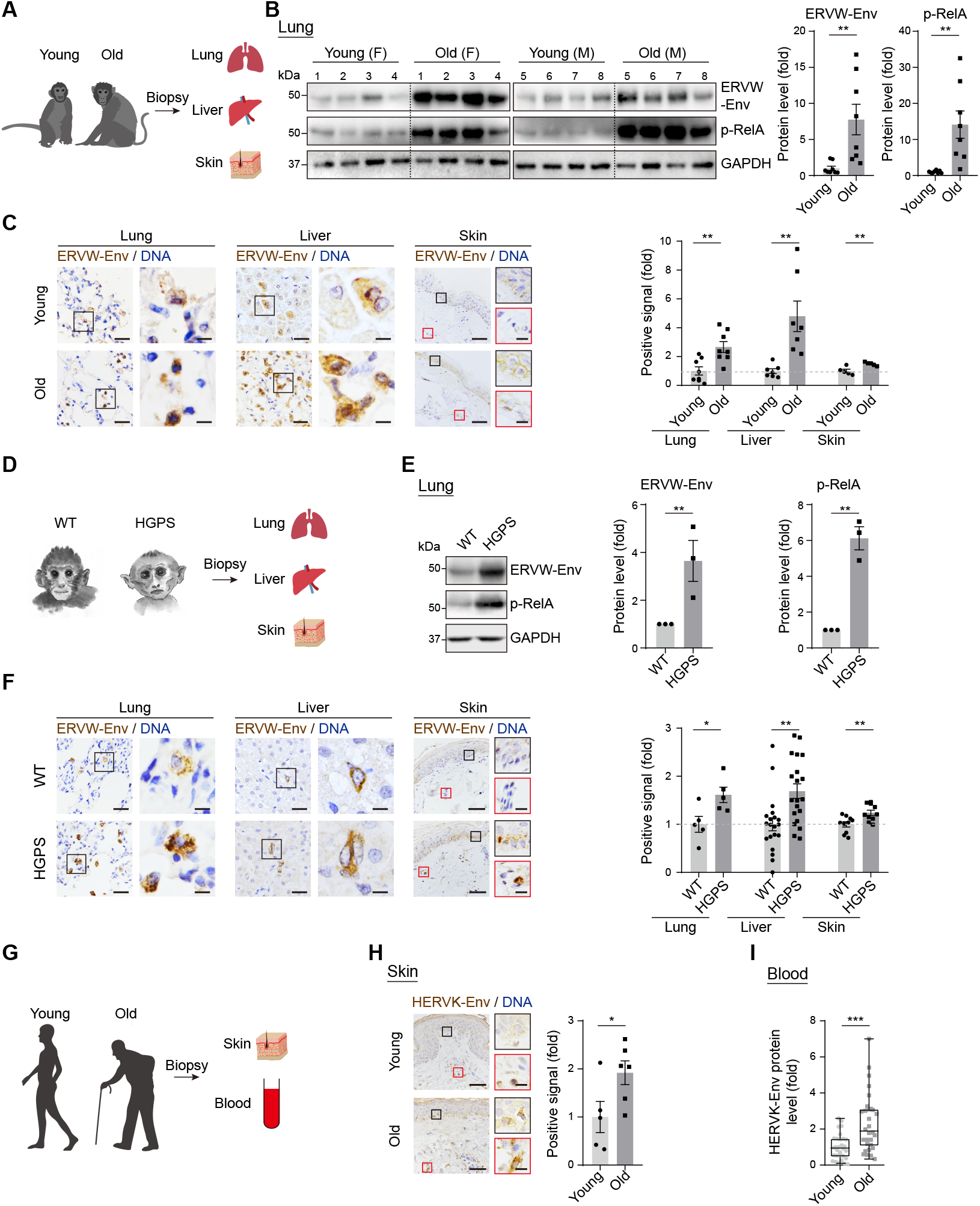
Activation of endogenous retrovirus as a biomarker of aging. (A-F) Schematic diagram of samples (A), western blotting of ERVW-Env and p-RelA in the lungs (B), and immunohistochemistry analysis of ERVW-Env in the lungs, livers, and skin (C) from young and old cynomolgus monkeys, or those from WT and HGPS cynomolgus monkeys (D-F). (G-I) Schematic diagram of samples (G), immunohistochemistry analysis of HERVK-Env in the skins (H), and ELISA analysis of HERVK-Env levels in serum (I) from young and old human donors. Scale bars, 50 μm and 10 μm (zoomed-in image) (all panels). See also Figure S6 and Table S2.

Finally, we sought to investigate whether elevated expression of HERVK is observed in human tissues during physiological aging. Likewise, in skin and serum samples obtained from young and old donors,^63^ HERVK-Env expression was markedly increased with age (Figures 6G–6I; Table S2). To ascertain whether HERVK in the serum from old individuals is a factor that drives aging, we cultured young primary hMPCs with medium that contained serum from young or old individuals (Figure S6E). Strikingly, we found that serum from old individuals increased HERVK abundance, elicited innate immune responses, and accelerated cellular senescence in primary hMPCs, while this pro-senescence capability was abolished upon HERVK immunodepletion (Figures S6F–S6H). Taken together, these results indicate that human endogenous retrovirus HERVK may serve as a potential biomarker to assess human aging as well as a potential therapeutic target for alleviating tissue and cellular senescence.

### Targeting endogenous retrovirus alleviates tissue aging

Next, we attempted to inhibit the expression of endogenous retroviruses in aged mice. Unlike humans, mice harbor a range of active endogenous retroviruses, among which mouse mammary tumor virus (MMTV), is also a known beta retrovirus, and is mostly closely related to that of HERVK (HML-2).^64,65^ Similar to the increased levels of ERV elements as we observed in primate and human tissues during aging, MMTV-Env levels were increased in lung, liver, and skin tissues of aged mice relative to young mice, alongside activation of the innate immunity and inflammatory pathways therein (Figures S7A–S7C). Thus, with some evolutionary conservation, both primate and rodent endogenous retroviruses are reactivated during aging.

Articular degeneration, or osteoarthritis (OA), is the most common joint pathology with aging, and has been attributed to the senescence of mesenchymal progenitor cells.^66–72^ Its pathology is demarcated by decreased numbers of Ki67-positive cells and reduced cartilage thickness (Figures S7D and S7E). In aged mice, we found that MMTV was substantially upregulated in the articular cartilage (Figure 7A), suggesting that increased levels of endogenous retroviruses could be a potential driver for aging-associated articular degeneration. Consistently, when we performed intra-articular injections of lentivirus carrying CRISPRi-dCas9/sgMMTV (sgRNA targeting MMTV) to repress MMTV (Figures 7B, 7C, and S7F), we detected phenotypes indicative of alleviation of tissue aging (Figures 7D, 7E, S7G, and S7H). We also observed structural and functional improvements in the joints of aged mice upon MMTV inhibition, as revealed by increased cartilage thickness and joint bone density, as well as enhanced grip strength (Figures 7F, 7G, and S7I).

**Figure 7.**
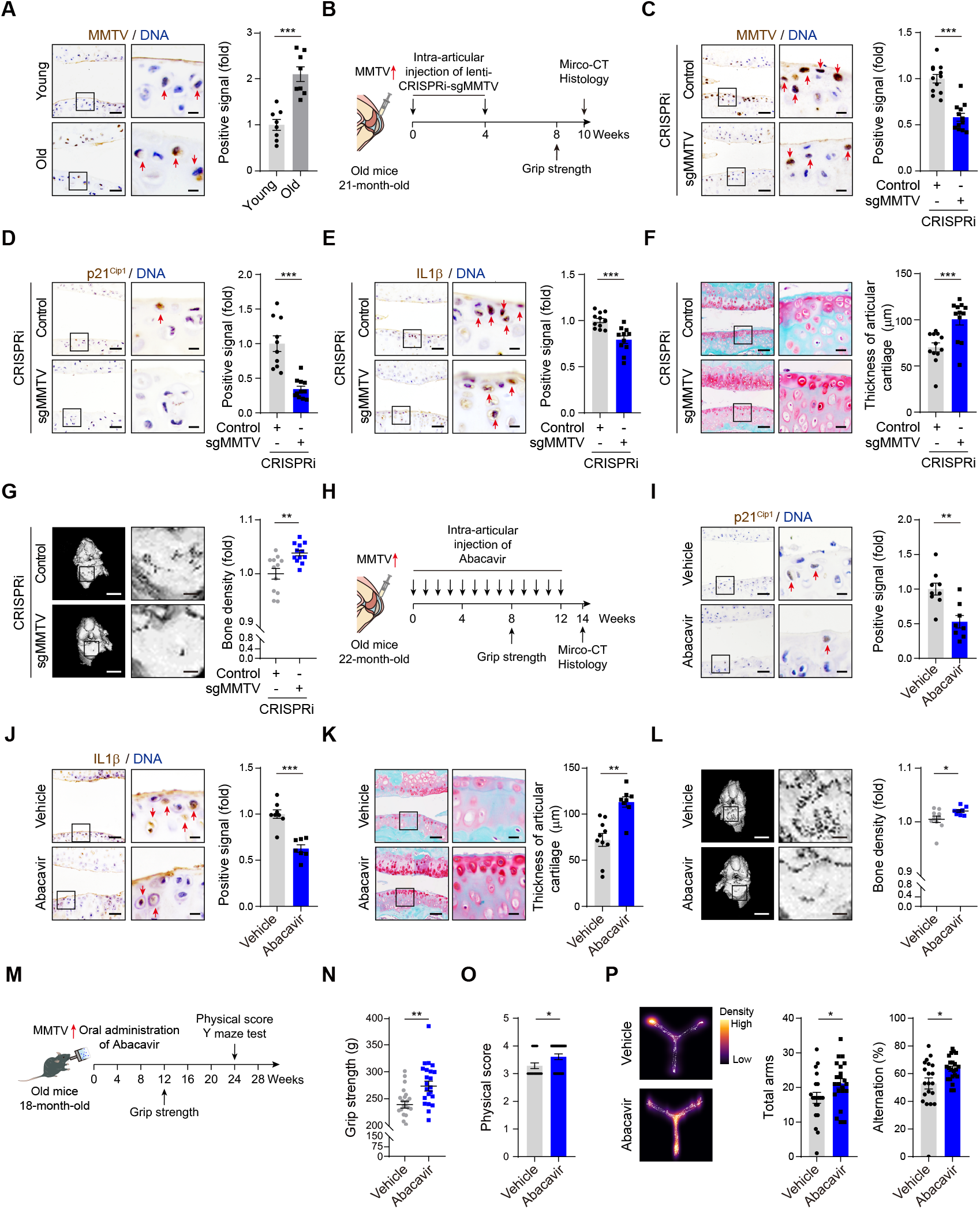
Targeting endogenous retrovirus alleviates tissue aging. (A) Immunohistochemistry analysis of MMTV-Env in the articular cartilages of young and old mice. (B-L) Schematic diagram showing the experimental procedure (B), immunohistochemistry analysis of MMTV-Env, p21^Cip1^, IL1β, and Safranin-O/Fast Green staining of the articular cartilages (C-F), as well as Micro-CT analysis showing the bone densities of the joints (G) of mice injected with lentiviruses expressing control or sgMMTV into articular cavities using a CRISPRi system, or those of mice intraarticularly injected with vehicle or Abacavir (H-L). (M-P) Schematic diagram showing the experimental procedure (M), grip strength analysis (N), overall physical scores (O), and Y maze analysis (P) of old mice fed with vehicle or Abacavir in the drinking water. Scale bars: 50 μm and 10 μm (zoomed-in images) in (A), (C)-(F), and (I)-(K); 500 μm and 100 μm (zoomed-in images) in (G) and (L). See also Figure S7.

Our findings that Abacavir treatment can inhibit the senescence-promoting effect of endogenous retrovirus (Figures 4N, 4O, 5P, 5S, S4K, and S5R) lay a foundation for further applying Abacavir to aging intervention *in vivo*. To this end, we performed weekly Abacavir injections into the articular cavities of 22-month-old mice (Figure 7H). Similar to lentiviral knockdown of MMTV using a CRISPRi system, we found that Abacavir treatment reduced cartilage degeneration, as evidenced by mitigated senescence and attenuated aging-associated inflammation (Figures 7I, 7J, S7J, and S7K). We also detected alleviation of aging-related articular degeneration, including augmented cartilage thickness, bone density, and grip strength (Figures 7K, 7L, and S7L). Finally, we asked whether Abacavir treatment could more generally improve health in aged mice. To this end, we treated 18-month-old mice with Abacavir dissolved in daily drinking water for 6 months (Figure 7M). Strikingly, we detected increased grip strength, improved overall physical score, and improved short-term memory in aged mice that were treated with Abacavir relative to untreated mice (Figures 7N–7P). Taken together, these data indicate that repression of endogenous retrovirus *in vivo* alleviates tissue aging and, to some extent, organismal aging.

## DISCUSSION

In this study, using various primate and rodent aging models, we discovered a positive feedback loop between endogenous retrovirus activation and aging. Our comprehensive analysis unraveled the causal relationship between HERVK and aging in multiple species, and was supported by pioneering studies showing increased expression of ERVs concomitant with aging in yeast, fly, and rodent models.^73–75^ For instance, activation of gypsy, a transposable element in Drosophila showing homology with vertebrate ERVs, was reported in aged flies.^76^ In addition, emerging studies suggested a correlation between awakened ERVs and aging-related disorders, such as rheumatoid arthritis and neurodegenerative diseases.^42,77–81^ More importantly, we successfully employed multiple strategies to block the pro-senescence effect of endogenous retroviruses, demonstrating the alleviation of aging defects across cellular models and multiple tissues *in vivo*. In line with our results, attempts to alleviate neurodegenerative disorders including amyotrophic lateral sclerosis (ALS), via inhibition of HERVK were reported.^82,83^ Thus, endogenous retroviruses represent druggable targets for alleviating aspects of aging and improving overall organismal health.

As another type of retrotransposon elements, LINE1 can also be activated during senescence and age-associated degeneration, and exert certain pro-senescence effects.^8,10,12,84,85^ Moreover, inhibition of LINE1 by reverse transcriptase inhibitors has been reported to alleviate aging-related phenotypes and extend the healthspan of mice.^9,11^ However, unlike LINE1, which is incapable of producing viral particles and thus acts primarily in a cell-autonomous manner, our study provides evidence that <u>a</u>ging-<u>i</u>nduced <u>r</u>esurrection of <u>e</u>ndogenous <u>r</u>etro<u>v</u>irus (AIR-ERV) not only exerts a devastating cell-autonomous role but also triggers secondary senescence in a paracrine manner.

In summary, our research provides experimental evidence that the conserved activation of endogenous retroviruses is a hallmark and driving force of cellular senescence and tissue aging. Our findings make fresh insights into understanding aging mechanisms, and especially enrich the theory of programmed aging. As such, our work lays the foundation for understanding the contagiousness of aging, opening avenues for establishing a scientific method for evaluating aging and developing clinical strategies to alleviate aging.

### Limitations of the study

Although we have provided evidence indicating the transmission process of HERVK RVLPs from senescent cells to young ones, more advanced technologies are required to monitor the exocytosis and endocytosis of viral particles by the donor and recipient cells in real time. It should be noted that a careful decision based on multiple indicators, beyond SA-β-gal activity alone,^86,87^ is urgently needed when evaluating HERVK-mediated cellular senescence. In addition, as short-read sequencing cannot be utilized to pinpoint the full-length copy of HERVK, further investigations are necessary to address whether an intact HERVK provirus may exist and how the full lifecycle of a HERVK provirus is completed during aging. As we demonstrated the effectiveness of aging alleviation via targeted inhibition of HERVK/MMTV in hMPCs and mice, it will be of great significance to perform preclinical trials in non-human primates for future translational applications. In addition, future studies will be needed to characterize endogenous retrovirus expression profiles across ages and species in large populations and examine the results with other aging clocks, such as DNA methylation and telomere length.

## STAR ★ METHODS

Detailed methods are provided in the online version of this paper and include the following:

- KEY RESOURCES TABLE
- RESOURCES AVAILABILITY
  - **Lead Contact**
  - **Materials Availability**
  - **Data and Code Availability**
- EXPERIMENTAL MODELS AND SUBJECT DETAILS
  - **Animals and human samples and ethics**
  - **Cell culture**
- METHOD DETAILS
  - **Animal experiments**
  - **Grip strength test**
  - **Y maze test**
  - **Micro-Computed Tomography (Micro-CT)**
  - **Western blotting**
  - **Immunoprecipitation (IP) of HERVK from conditioned medium**
  - **Purification of microvesicle particles from conditioned medium**
  - **DNA/RNA isolation and (reverse transcription-) quantitative PCR (q(RT-)PCR))**
  - **Viral RNA isolation from microvesicles**
  - **Droplet Digital PCR (ddPCR)**
  - **ELISA**
  - **Immunofluorescence, immunohistochemistry staining and microscopy**
  - **(sm)RNA/DNA-FISH**
  - **(Immuno-)Transmission electron microscopy (TEM)**
  - **Plasmid construction**
  - **HERVK RVLPs and lentivirus production**
  - **Cell treatment**
  - **Clonal expansion assay**
  - **SA-β-gal staining**
  - **ChIP-qPCR**
  - **Strand-specific RNA sequencing (RNA-seq)**
  - **RNA-seq data processing**
  - **Analysis of the expression levels of repetitive elements**
  - **Whole genome bisulfite sequencing (WGBS) library construction and sequencing**
  - **WGBS data processing**
  - **Quantification and statistical analysis**

## SUPPLEMENTAL INFORMATION

Supplemental information can be found online at http://…

## ACKNOWLEDGEMENTS

We would like to thank the Center for Biological Imaging (CBI), Institute of Biophysics (Chinese Academy of Science) for TEM analysis, and we would be grateful to C. Peng for TEM sample preparation and L. Zhang for the help with the TEM operation. We would also like to express our gratitude to Y. Li (Tsinghua University) for immuno-TEM sample preparation. We thank Z. Zhang and Y. Tao (Spatial FISH, Co., Ltd) for the technical help in the FISH experiment, as well as S. Li and J. Hao (the Center of Imaging facility of Institute of Zoology and Institute of Biophysics (Chinese Academy of Sciences), for the help with imaging scanning. We would also like to thank Dr. G. Pei for his insightful comments. We are grateful to J. Jia, L. Huang and W. Wang for their help in animal experiments, as well as L. Bai, Q. Chu, L. Tian, J. Lu, Y. Yang, X. Li, J. Chen, R. Bai, and X. Jing for administrative assistance. We also would like to thank the Chinese Primate Biomedical Research Alliance (CPBRA), and the Aging Biomarker Consortium, China (ABC) for supporting our study. This work was supported by the National Key R&D Program of China (2020YFA0804000), the Strategic Priority Research Program of the Chinese Academy of Sciences (XDA16000000), the National Key R&D Program of China (2022YFA1103700, 2018YFC2000100, 2018YFA0107203, 2020YFA0112200, 2019YFA0802202, 2020YFA0803401, 2021YFF1201005, 2022YFA1103800, and 2019YFA0110100), the National Natural Science Foundation of China (81921006, 82125011, 92149301, 92168201, 91949209, 92049304, 92049116, 32121001, 82192863, 82122024, 82071588, 31970597, 81861168034, 31900524, 32100937, 32000510, 32200610, 82201727), the STI2030-Major Projects (2021ZD0202400), CAS Project for Young Scientists in Basic Research (YSBR-076, YSBR-012), the Program of the Beijing Natural Science Foundation (Z190019), K. C. Wong Education Foundation (GJTD-2019-06, GJTD-2019-08), The Pilot Project for Public Welfare Development and Reform of Beijing-affiliated Medical Research Institutes (11000022T000000461062), Youth Innovation Promotion Association of CAS (E1CAZW0401), Young Elite Scientists Sponsorship Program by CAST (YESS20200012), the Informatization Plan of Chinese Academy of Sciences (CAS-WX2021SF-0301, CAS-WX2022SDC-XK14, CAS-WX2021SF-0101), and the Tencent Foundation (2021-1045).

## AUTHOR CONTRIBUTIONS

G.-H.L., J.Q., W.Z. conceptualized the work and supervised the overall experiments. X.L. performed the cell culture, and the experiments related to functional and mechanistic analyses, including western blotting, qRT-PCR, ddPCR, ChIP-qPCR and immunofluorescence staining. Z.L. performed bioinformatic analyses. Z.W. performed cell culture and sequencing library construction. J.R. performed the manuscript writing. L.S. provided human serum samples. Y.F. performed immunohistochemistry staining in cynomolgus monkeys and humans. Y.N. provided HGPS cynomolgus monkey samples. G.C. and L.L. performed smRNA/DNA-FISH. X.J. performed fibroblast cell culture and functional analyses. Q.J. Y.E.Z., S.T., Y.C. helped with the bioinformatic analyses. B.Z. performed statistical analysis of immunohistochemistry staining. C.W. performed the cell culture. Q.W. helped with the schematic diagram drawing. K.Y. helped with vector constructions. W.L. organized the animal experiments. A.Z. helped with virus purification. W.J., Q.Z., J.C.I.B. C.R.E., F.L., and F.T.helped with supervision of the project. G.-H.L., J.Q., W.Z., J.R., X.L., Z.L., Z.W., Z.J., Y.S.C., S.W., and M.S. performed manuscript writing, reviewing, and editing. All authors reviewed the manuscript.

## Supplemental Figure Legends

**Figure S1.**
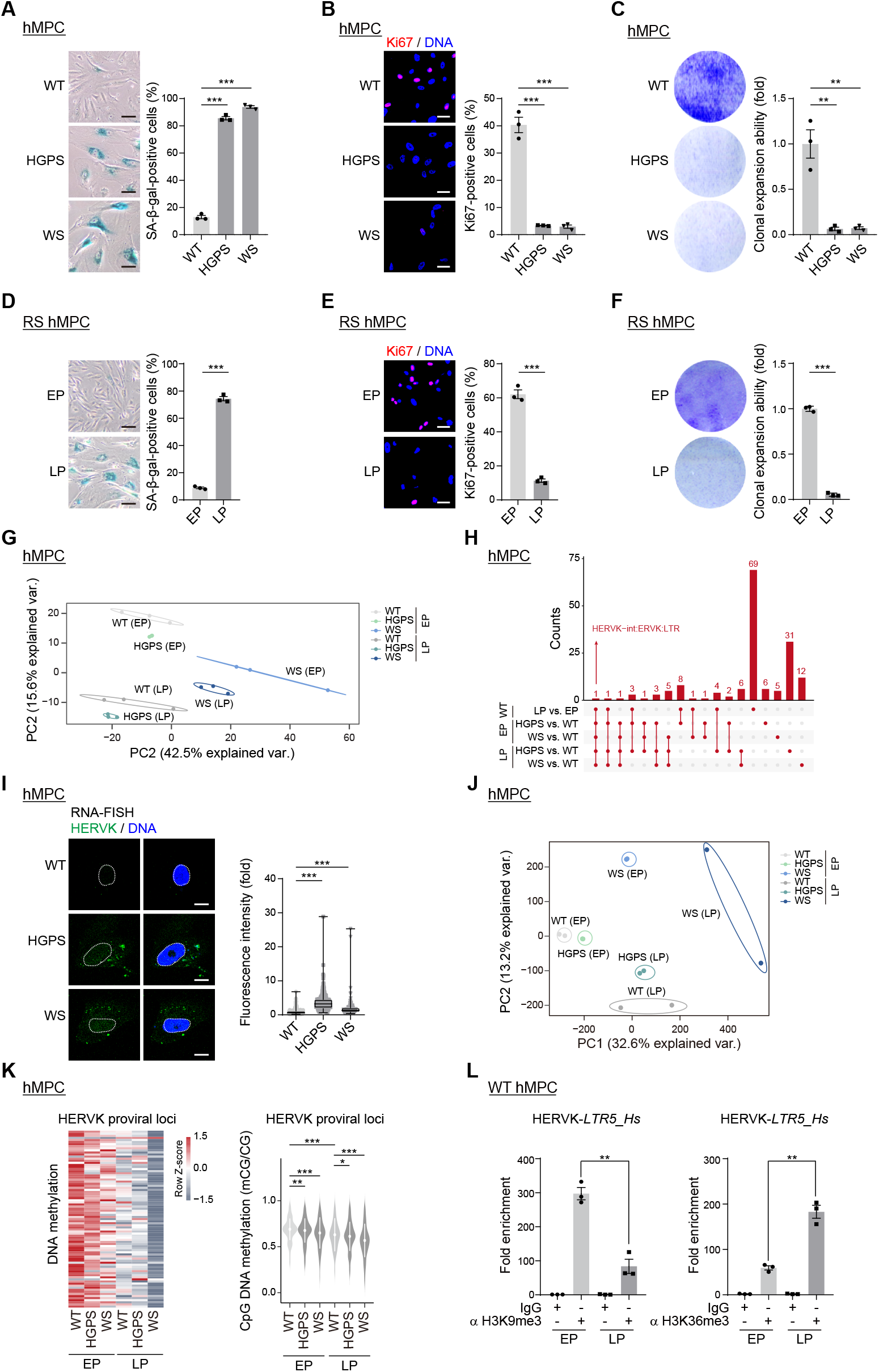
Increased expression of HERVK in senescent hMPCs, related to Figure 1. (A-F) SA-β-gal, Ki67 staining, and clonal expansion assay of WT, HGPS and WS hMPCs, as well as WT hMPCs at EP and LP. (G-H) Principal component analysis (PCA) of RNA-seq data based on the normalized read count of repetitive elements (G), and UpSet plot showing the overlap of upregulated RepeatMasker-annotated repetitive elements (H) in RS and prematurely senescent hMPCs. (I) HERVK RNA-FISH analysis in WT, HGPS and WS hMPCs. (J-K) PCA of the whole-genome CpG DNA methylation levels (J) and CpG methylation levels at the HERVK proviral genomic locus (K) in RS and prematurely senescent hMPCs. (L) ChIP-qPCR analysis of H3K9me3 and H3K36me3 enrichment in HERVK-*LTR5_Hs* regions in RS hMPCs. Scale bars: 20 μm in (A), (B), (D), and (E); and 10 μm in (I).

**Figure S2.**
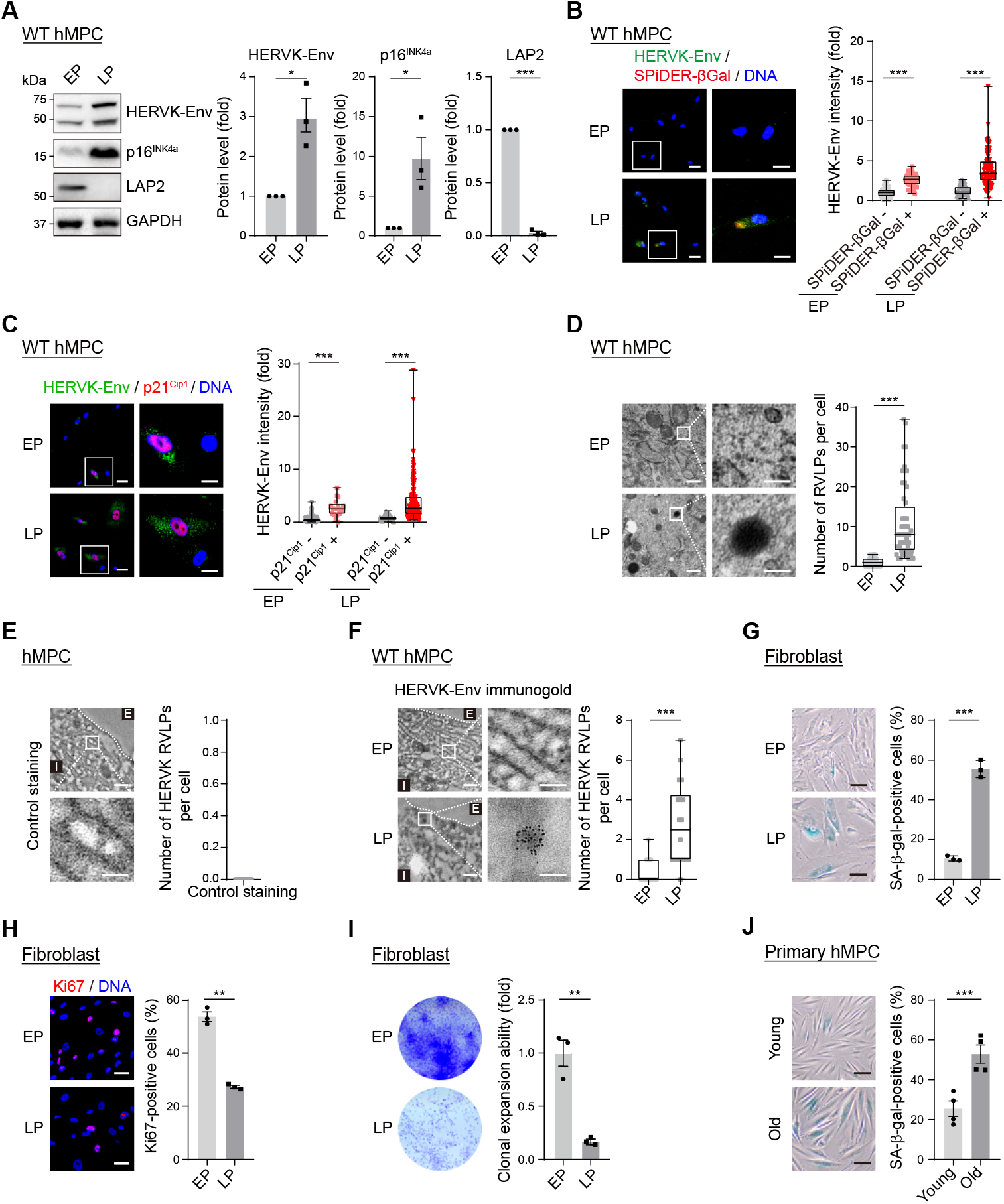
Increased HERVK proviral components in senescent cells, related to Figure 2. (A-C) Western blotting of HERVK-Env, p16^INK4a^ and LAP2 (A), immunofluorescence co-staining of HERVK-Env and SPiDER-βGal (B) or p21^Cip1^ (C) in RS WT hMPCs. (D-F) TEM analysis (D), and TEM analysis after negative immunogold labeling without the primary antibody (E), or immunogold labeling with anti-HERVK-Env antibody (F) in RS WT hMPCs. (G-J) SA-β-gal, Ki67 staining, and clonal expansion assay of RS human fibroblasts (G-I), as well as SA-β-gal staining of primary hMPCs from young and old donors (J). Scale bars: 20 μm and 10 μm (zoomed-in images) in (B) and (C); 200 nm and 100 nm (zoomed-in images) in (D)-(F); and 20 μm in (G), (H), and (J).

**Figure S3.**
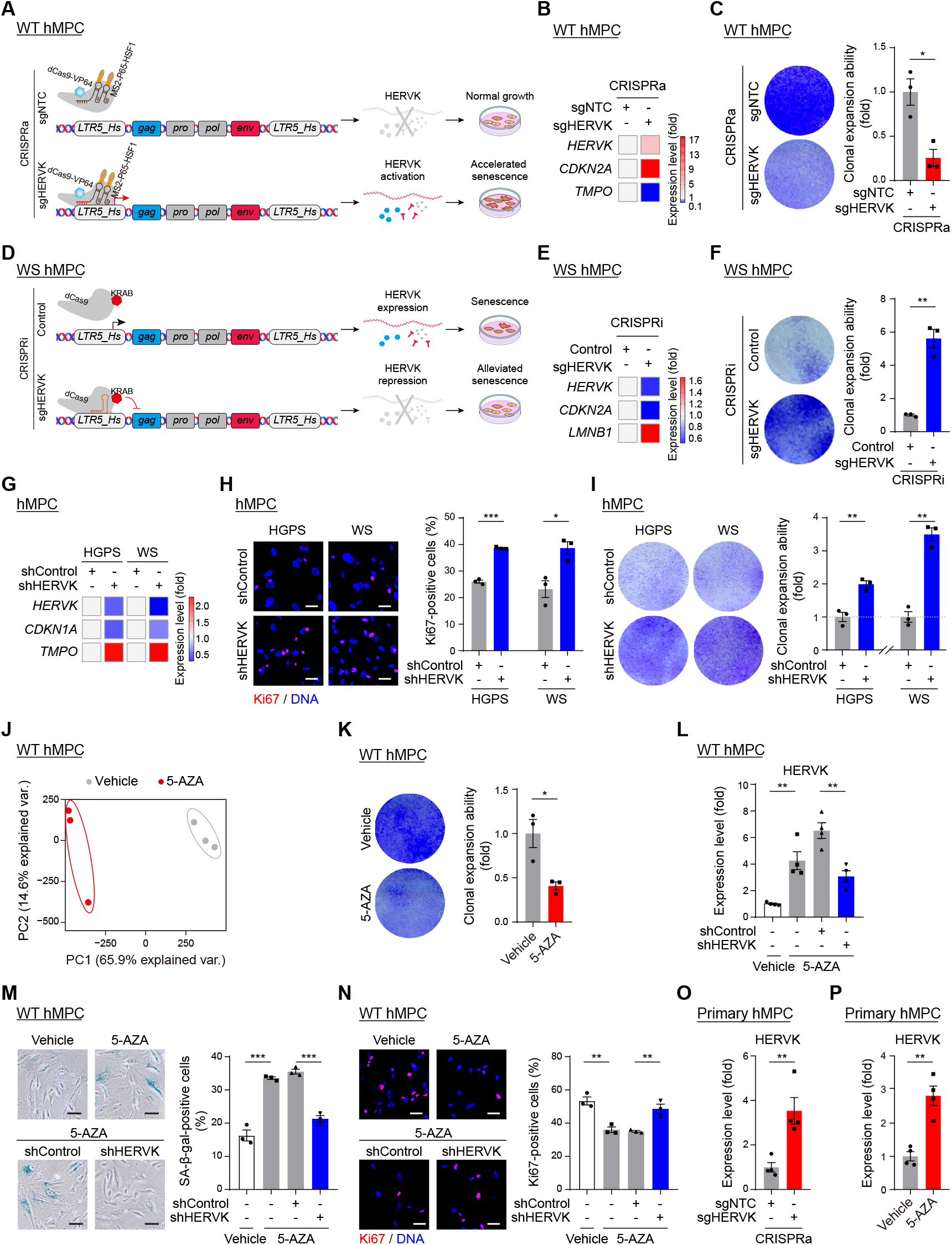
The upregulation of HERVK drives hMPC senescence, related to Figure 3. (A-F) Schematic diagram of the experimental procedure (A), heatmap showing the qRT-PCR analysis of HERVK and senescence marker genes (B), and clonal expansion assay (C) in WT hMPCs transduced with lentiviruses expressing sgNTC or sgHERVK using a CRISPRa system, or those in WS hMPCs transduced with lentiviruses expressing control or sgHERVK using a CRISPRi system (D-F). (G-I) Heatmap showing the qRT-PCR analysis of HERVK and senescence marker genes (G), Ki67 staining (H) and clonal expansion assay (I) of HGPS or WS hMPCs after transduction with lentivirus delivering shControl or shHERVK. (J-K) PCA of the whole-genome CpG DNA methylation levels (J) and clonal expansion assay (K) in vehicle- or 5-AZA-treated WT hMPCs. (L-N) qRT-PCR analysis of the HERVK levels (L), SA-β-gal (M) and Ki67 staining (N) of vehicle- or 5-AZA-treated WT hMPCs after HERVK knockdown. (O-P) qRT-PCR analysis of the HERVK levels in primary hMPCs from a young individual transduced with lentiviruses expressing sgNTC or sgHERVK using a CRISPRa system (O) or that treated with vehicle or 5-AZA (P). Scale bars, 20 μm (all panels).

**Figure S4.**
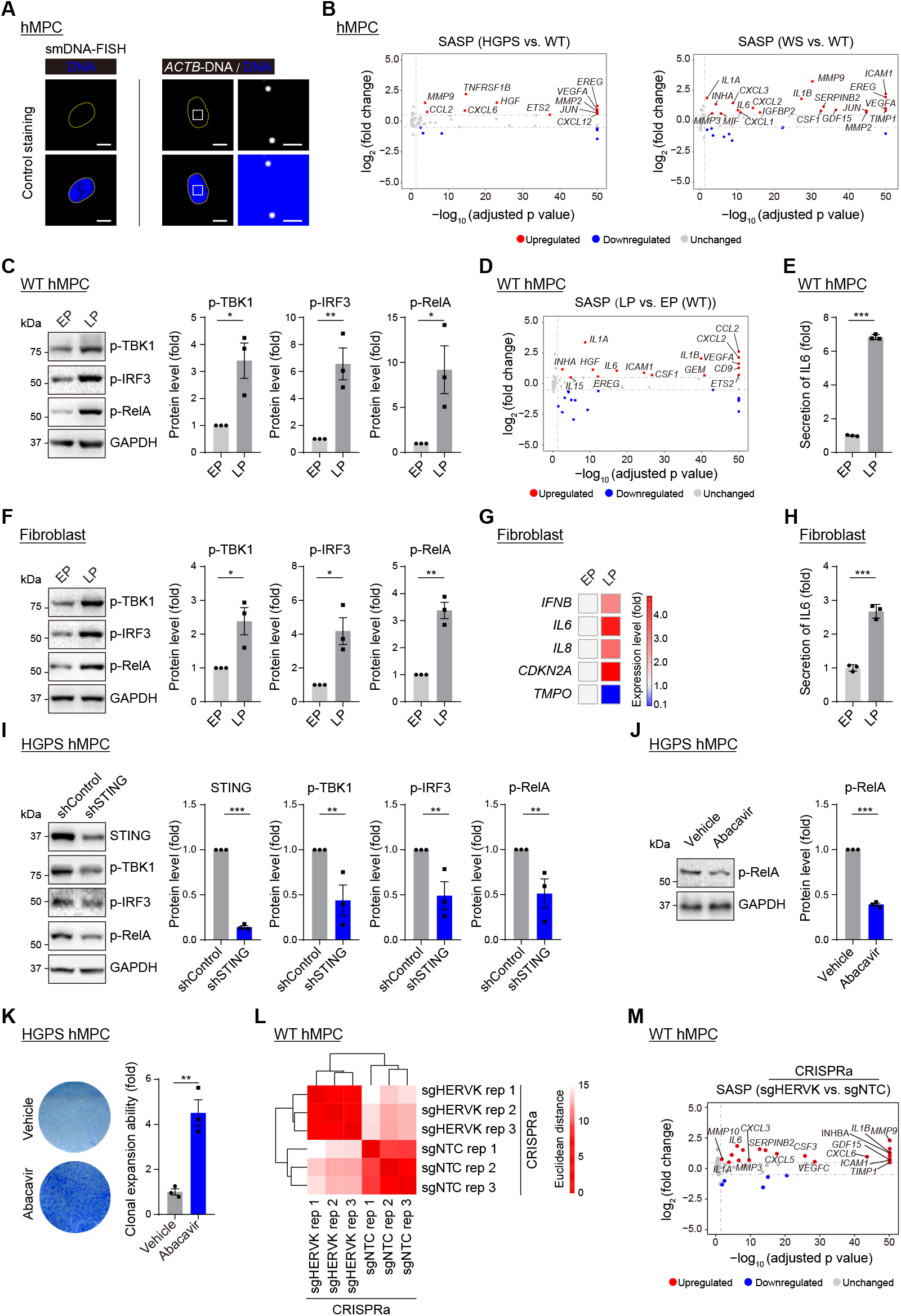
The innate immune response is activated in senescent cells, related to Figure 4. (A) Negative and positive control staining (*ACTB*) for smDNA-FISH in hMPCs. (B) Scatter plots showing the differential expression levels of SASP genes in HGPS or WS hMPCs compared to WT hMPCs at LP. (C-E) Western blotting of p-TBK1, p-IRF3 and p-RelA (C), as well as scatter plot showing the differential expression levels of SASP genes (D) and ELISA analysis of IL6 levels in the culture medium (E), in RS WT hMPCs. (F-H) Western blotting of p-TBK1, p-IRF3 and p-RelA (F), as well as qRT-PCR analysis of SASP genes (G) and ELISA analysis of IL6 levels in culture medium (H) in RS fibroblasts. (I) Western blotting of STING, p-TBK1, p-IRF3 and p-RelA in HGPS hMPCs after STING knockdown. (J-K) Western blotting of p-RelA (J) and clonal expansion assay (K) in HGPS hMPCs treated with Abacavir. (L-M) Heatmap showing the Euclidean distance analysis to evaluate the reproducibility of the RNA-seq (L) and scatter plots showing the differential expression levels of SASP genes (M) in WT hMPCs transduced with the lentivirus expressing sgNTC or sgHERVK using the CRISPRa system. Scale bars: 10 μm and 200 nm (zoomed-in images) in (A).

**Figure S5.**
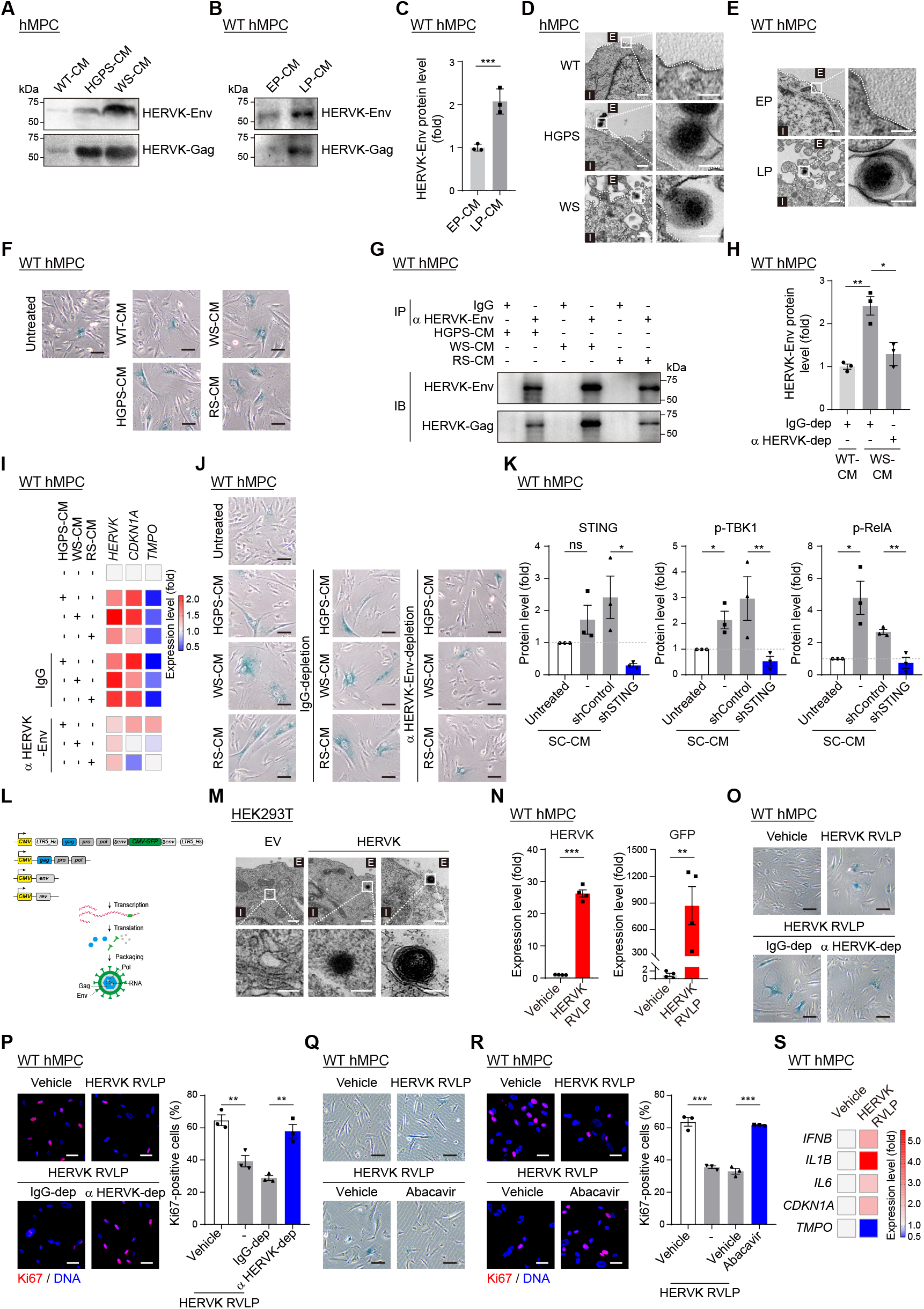
HERVK released from senescent cells induces young cell senescence, related to Figure 5. (A-B) Western blotting of HERVK-Env and -Gag in microvesicles from CM of WT, HGPS and WS hMPCs (A), as well as RS WT hMPCs (B). (C) ELISA analysis of relative HERVK-Env levels in the CM of RS WT hMPCs. (D-E) TEM images of WT, HGPS and WS hMPCs (D), as well as RS WT hMPCs (E). (F) Representative images of SA-β-gal staining for Figure 5G. (G-J) Western blotting of HERVK-Env and -Gag in the immunoprecipitates of SC-CM (G), ELISA analysis of relative HERVK-Env protein levels in the SC-CM (H), and heatmap showing qRT-PCR analysis of the HERVK levels (I) and representative images of SA-β-gal staining for Figure 5I (J) in WT hMPCs treated with SC-CM, after immunodepletion with IgG or anti-HERVK antibody. (K) Statistical results for western blotting of STING, p-TBK1 and p-RelA in Figure 5J. (L-M) Schematic diagram showing the constructs to produce HERVK RVLP (L), and TEM images of HEK293T cells transfected with empty vector (EV) or HERVK constructs (M). (N-O) qRT-PCR analysis showing the levels of HERVK and GFP (N), representative images of SA-β-gal staining for Figure 5O (O) in WT hMPCs infected with HERVK RVLPs. (P) Immunofluorescence staining of Ki67 in WT hMPCs infected with HERVK RVLPs after pretreatment with IgG or anti-HERVK-Env antibody. (Q-R) Representative images of SA-β-gal staining for Figure 5P (Q) and immunofluorescence staining of Ki67 (R) in WT hMPCs infected with HERVK RVLPs followed by Abacavir treatment (S) Heatmap showing the qRT-PCR analysis of the expression levels of SASP genes and senescence marker genes in WT hMPCs treated with vehicle or HERVK RVLPs. Scale bars: 200 nm and 100 nm (zoomed-in images) in (D), (E), and (M); 20 μm in (F), (J), and (O)-(R).

**Figure S6.**
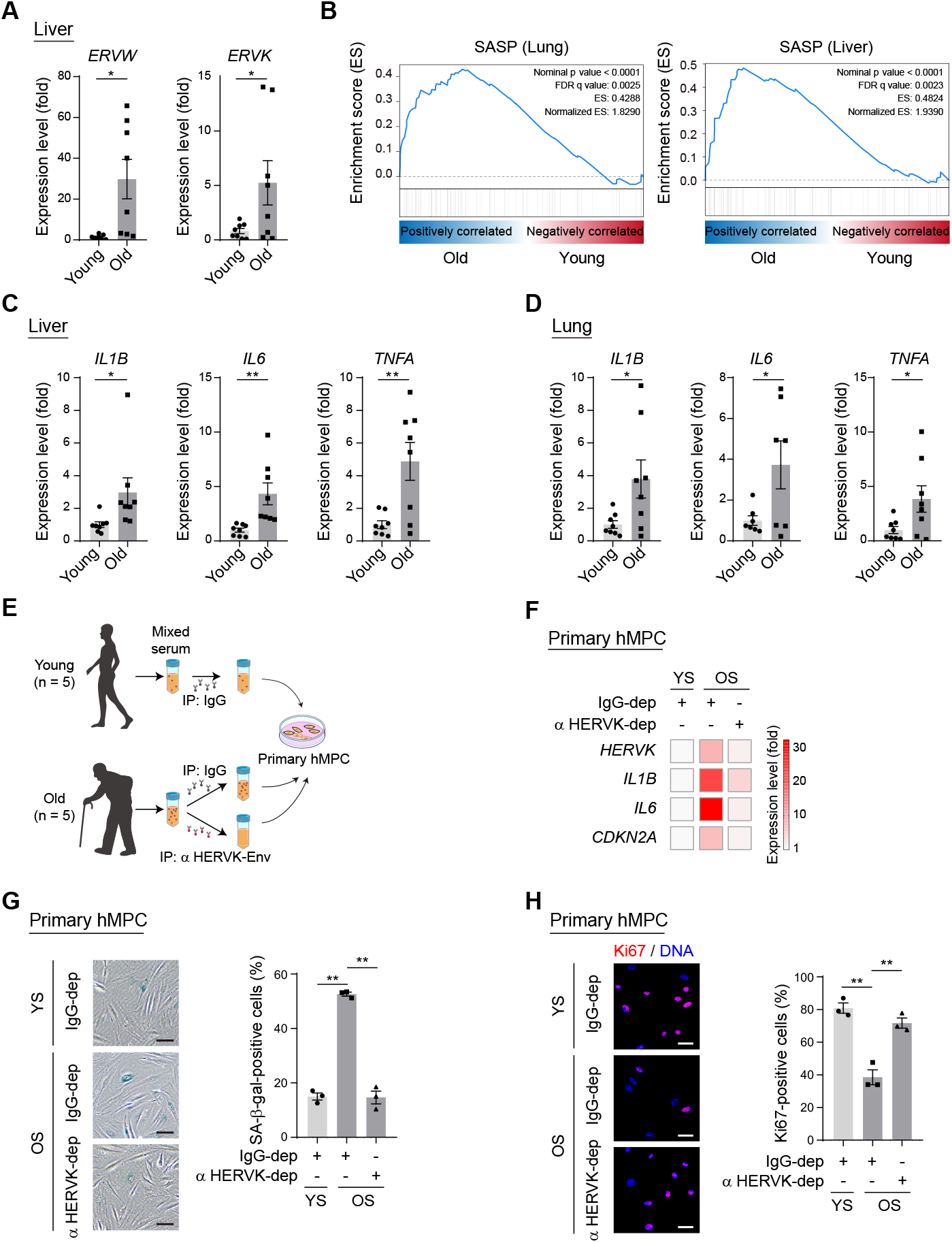
Activation of endogenous retrovirus and the innate immune pathway *in vivo*, related to Figure 6. (A-D) qRT-PCR analysis of the levels of ERVW and ERVK in the livers (A), gene set enrichment analysis (GSEA) of SASP gene levels in lungs and livers (B), and qRT-PCR analysis of the levels of SASP genes in the livers (C) and lungs (D) of young and old cynomolgus monkeys. (E-H) Schematic diagram showing the experimental procedure (E), qRT-PCR analysis of the expression levels of HERVK, SASP genes and senescence marker genes (F), as well as SA-β-gal (G) and Ki67 (H) staining of young primary hMPCs cultured with medium containing young (YS) or old sera (OS) after immunodepletion with using IgG or anti-HERVK antibody. Scale bars, 20 μm (all panels).

**Figure S7.**
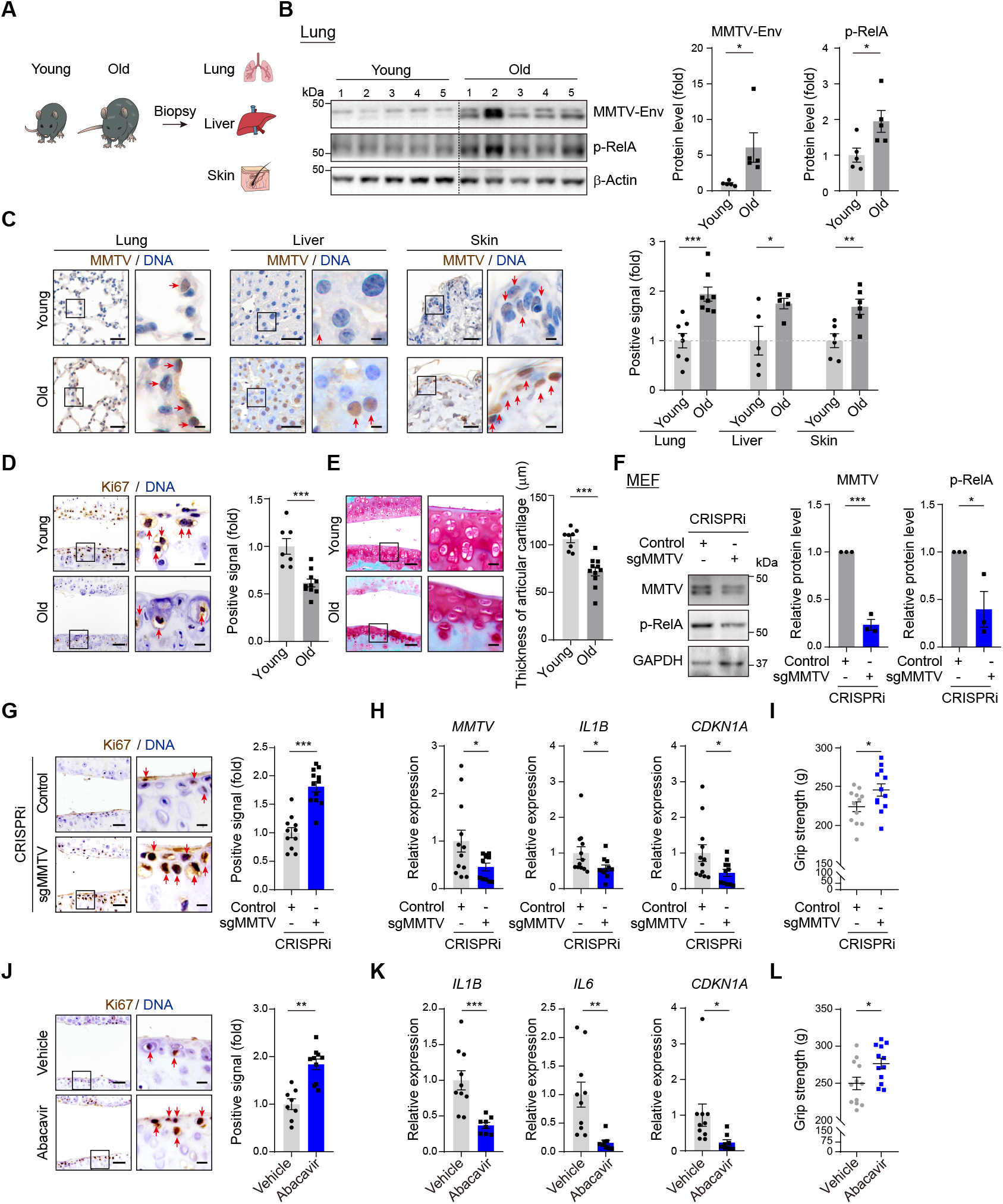
Suppression of endogenous retrovirus alleviates tissue aging in mice, related to Figure 7. (A-C) Schematic diagram of samples (A), western blotting of MMTV and p-RelA in the lungs (B), and immunohistochemistry analysis of MMTV in the lungs, livers, and skin (C), from young and old mice, (D-E) Immunohistochemistry analysis of Ki67 (D) and Safranin-O/Fast Green (E) staining in the articular cartilages of young and old mice. (F) Western blotting of MMTV and p-RelA in MEFs transduced with lentiviruses expressing control or sgMMTV using a CRISPRi system. (G-I) Immunohistochemistry analysis of Ki67 (G), qRT-PCR analysis of the levels of *MMTV, IL1B* and *CDKN1A* in the articular cartilages (H), and grip strength analysis (I) of mice intra-articularly injected with lentiviruses expressing control or sgMMTV using a CRISPRi system. (J-L) Immunohistochemistry analysis of Ki67 (J), qRT-PCR analysis of the levels of *IL1B, IL6* and *CDKN1A* levels in the articular cartilages (K), and grip strength analysis (L) of mice injected with vehicle or Abacavir. Scale bars, 50 μm and 10 μm (zoomed-in images) (all panels).

## STAR ★ METHODS

### KEY RESOURCES TABLE

| Antibodies |  |  |
| --- | --- | --- |
| Mouse monoclonal anti-HERVK-Env | Austral Biologicals | Cat# HERM-1811-5;<br>RRID:<br>AB_10891119 |
| Mouse monoclonal anti-HERVK-Gag | Austral Biologicals | Cat# HERM-1841-5;<br>RRID:<br>AB_10891407 |
| Rabbit polyclonal anti-ERVK7 | United States Biological | Cat# 302427<br>RRID: N/A |
| Rabbit polyclonal anti-ERVW-1 | Abcam | Cat# ab179693;<br>RRID: N/A |
| Rabbit anti-phospho-TBK1/NAK (Ser172) | Cell Signaling Technology | Cat# 5483;<br>RRID:<br>AB_10693472 |
| Rabbit anti-phospho-NF-κB p65 (p-RelA) (Ser536) (93H1) | Cell Signaling Technology | Cat# 3033S;<br>RRID: AB_331284 |
| Rabbit anti-phospho-IRF-3 (Ser396) | Cell Signaling Technology | Cat# 29047S;<br>RRID: AB_2773013 |
| Mouse monoclonal anti-STING | R&D systems | Cat#MAB7169;<br>RRID:<br>AB_10971940 |
| Mouse monoclonal anti-p16<sup>INK4a</sup> | BD Biosciences | Cat# 550834;<br>RRID: AB_2078446 |
| Mouse anti-LAP2 | BD Biosciences | Cat# 611000;<br>RRID: AB_398313 |
| Rabbit monoclonal anti-p21<sup>Cip1</sup> | Cell Signaling Technology | Cat# 2947s;<br>RRID: AB_823586 |
| Rabbit monoclonal anti-p21<sup>Cip1</sup> | Abcam | Cat# ab188224;<br>RRID: AB_2734729 |
| Mouse monoclonal anti-GAPDH | Santa Cruz Biotechnology | Cat#; sc-365062;<br>RRID:<br>AB_10847862 |
| Mouse monoclonal anti-β-Actin | Santa Cruz Biotechnology | Cat# sc-69879;<br>RRID: AB_1119529 |
| Rabbit polyclonal anti-H3K9me3 | Abcam | Cat# ab8898;<br>RRID: AB_1119529 |
| Rabbit polyclonal anti-H3K36me3 | Abcam | Cat# ab9050;<br>RRID: AB_306966 |
| Mouse monoclonal anti-Ki67 | ZSGB-Bio | Cat# ZM-0166;<br>RRID: AB_2890067 |
| Rabbit polyclonal anti-Ki67 | Abcam | Cat# ab15580;<br>RRID: AB_443209 |
| Rabbit monoclonal anti-cGAS (D1D3G) | Cell Signaling Technology | Cat# 15102S;<br>RRID: AB_2732795 |
| Rabbit polyclonal anti-MMTV-Env | Novus Biologicals, LLC | Cat# NBP2-44179;<br>RRID: N/A |
| Mouse monoclonal anti-IL1β | Santa Cruz Biotechnology | Cat# sc-52012;<br>RRID: AB_629741 |
| Normal Rabbit IgG | Cell Signaling Technology | Cat# 2729S;<br>RRID: AB_1031062 |
| Normal Mouse IgG | Santa Cruz Biotechnology | Cat# sc-2025;<br>RRID: AB_737182 |
| Alexa Fluor™ 568 Donkey Anti-Rabbit IgG (H+L) | Invitrogen | Cat# A10042;<br>RRID: AB_2534017 |
| Alexa Fluor™ 568 Donkey Anti-Mouse IgG (H+L) | Invitrogen | Cat# A-10037;<br>RRID: AB_2534013 |
| Alexa Fluor™ 488 Donkey Anti-Mouse IgG (H+L) | Invitrogen | Cat# A21202;<br>RRID: AB_141607 |
| HRP-conjugated Goat Anti-Mouse IgG (H+L) | ZSGB-Bio | Cat# ZB2305;<br>RRID: AB_2747415 |
| HRP-conjugated Goat Anti-Rabbit IgG (H+L) | ZSGB-Bio | Cat# ZB2301;<br>RRID: AB_2747412 |
| 6-nm Gold-labeled Goat Anti-Mouse IgG (H+L) | Jackson ImmunoResearch | Cat# 115-195-146<br>RRID: N/A |
| **Biological Samples** |  |  |
| The young and old cynomolgus monkey tissues | Xieerxin Biology Resource | N/A |
| The WT and HGPS cynomolgus monkey tissues | Yunnan Key Laboratory of Primate Biomedical Research | N/A |
| Human eyelid skin samples | Peking Union Medical College Hospital | N/A |
| Human serum samples | The First Hospital of Kunming Medical University | N/A |
| **Chemicals, Peptides, and Recombinant Proteins** |  |  |
| MEMα | GIBCO | Cat# 32571-101 |
| DMEM/High Glucose | HyClone | Cat# SH30243.01 |
| Non-essential amino acids (NEAA) | GIBCO | Cat# 11140076 |
| Fetal bovine serum (FBS) | GIBCO | Cat# 42Q7980K |
| Penicillin/streptomycin (P/S) | GIBCO | Cat# 15140-163 |
| GlutaMAX | GIBCO | Cat# 35050079 |
| Gelatin | Sigma-Aldrich | Cat# V900863 |
| bFGF | Joint Protein Central | Cat# BBI-EXP-002 |
| TrypLE Express Enzyme (1X), no phenol red | GIBCO | Cat# 12605010 |
| 0.25% Trypsin-EDTA (1X), Phenol Red | GIBCO | Cat# 25200072 |
| Hoechst 33342 | Thermo Fisher Scientific | Cat# H3570 |
| THUNDERBIRD SYBR qPCR mix | TOYOBO | Cat# QPS-201(-) |
| 4% paraformaldehyde, PFA | Dingguo, China | Cat# AR-0211 |
| Triton X-100 | Sigma-Aldrich | Cat# T9284 |
| Crystal violet | Biohao Biotechnology Co., Ltd., China | Cat# C0502 |
| TRIzol™ Reagent | Thermo Fisher Scientific | Cat# 15596018 |
| DMSO | Sigma-Aldrich | Cat# D2650 |
| NP-40 | Dingguo, China | Cat# DH218 |
| Proteinase K | New England Biolabs | Cat# P8107S |
| RNase A | TAKARA | Cat# 2158 |
| EDTA | Sigma-Aldrich | Cat# EDS-500G |
| Protease inhibitor cocktail | Roche | Cat# 4693159001 |
| X-gal | Amresca | Cat# 0428-25G |
| PVDF membranes | Millipore | Cat# IPVH00010 |
| Protein A/G Plus-Agarose | Thermo Fisher Scientific | Cat# sc-2003 |
| QX200 ddPCR EvaGreen Supermix | Bio-Rad Laboratories, Inc. | Cat# 186-4033 |
| QX200™ Droplet Generation Oil for EvaGreen | Bio-Rad Laboratories, Inc. | Cat# 1864005 |
| MluI | New England Biolabs | Cat# R0198S |
| ClaI | New England Biolabs | Cat# R0197S |
| ESP3I/BsmB1 | FERMENTAS | Cat# FD0454 |
| NheI | New England Biolabs | Cat# R3131S |
| ApaI | New England Biolabs | Cat# R0114S |
| KpnI-HF | New England Biolabs | Cat# R3142 |
| BamHI-HF | New England Biolabs | Cat# R3136 |
| 5-AZA | Sigma-Aldrich | Cat# 320-67-2 |
| Abacavir | Selleckchem | Cat# S3165 |
| Polybrene | Sigma-Aldrich | Cat# 107689 |
| **Critical Commercial Assays** |  |  |
| BCA Protein Assay Kit | Dingguo, China | Cat# BCA02 |
| Qubit™ dsDNA HS Assay Kit | Thermo Fisher Scientific | Cat# Q32854 |
| Dynabeads™ Protein A | Thermo Fisher Scientific | Cat# 10002D |
| Ribo-Zero™ rRNA Removal Kit | Epicentre Biotechnologies | Cat# RZH1046 |
| NEBNext® Ultra™ Directional RNA Library Prep Kit for Illumina® | New England Biolabs | Cat# E7420L |
| DNeasy Blood & Tissue Kit | QIAGEN | Cat# 69506 |
| GoScript Reverse Transcription System | Promega | Cat# A5001 |
| Lipofectamine 3000 Transfection Reagent | Thermo Fisher Scientific | Cat# L3000015 |
| QIAamp Viral RNA Mini Kit | QIAGEN | Cat# 52904 |
| SuperSignal™ West Femto Maximum Sensitivity Substrate | Thermo Fisher Scientific | Cat# 34096 |
| TURBO DNA-free™ Kit | Thermo Fisher Scientific | Cat# AM1907 |
| Human IL6 ELISA Kit | BioLegend | Cat# 430515 |
| Human HERVK-Env Polyprotein ELISA Kit | CUSABIO | Cat# CBS-EL007812HU |
| 2'3'-cGAMP ELISA Kit | Cayman | Cat# 501700 |
| DAB Staining Kit | ZSGB-BIO | Cat# ZLI-9018; PV-9001; PV-9002 |
| QuantiGene® ViewRNA ISH Cell Assay Kit | Thermo Fisher Scientific | Cat# QVC0001 |
| Cellular Senescence Detection Kit-SPiDER-βGal | DOJINDO Laboratories | Cat# PL507 |
| **Deposited Data** |  |  |
| The raw RNA-seq data for prematurely senescent and RS hMPCs | This study | CRA003282 |
| The raw WGBS data for prematurely senescent and RS hMPCs | Liu et al.<sup>39</sup> | HRA001144 |
| The raw WGBS data for vehicle or 5-AZA treated hMPCs | This study | HRA001879 |
| The raw RNA-seq data for WT MPCs (P6) after transduction of lentiviruses with sgRNA targeting HERVK or NTC using a CRISPRa system | This study | CRA003283 |
| The RNA-seq raw data liver tissues of young and old cynomolgus monkeys | Liu et al.<sup>66</sup> | CRA002965 |
| The RNA-seq raw data for lung tissues of young and old cynomolgus monkeys | Ma et al.<sup>88</sup> | CRA002810 |
| **Experimental Models: Cell Lines** |  |  |
| WT hMPCs (derived from WT hESCs (Line H9)) | Wu et al.<sup>33</sup> | N/A |
| *LMNA*<sup>G608G/+</sup> hMPCs | Wu et al.<sup>33</sup> | N/A |
| *WRN*<sup>-/-</sup> hMPCs | Zhang et al., Wu et al.<sup>32,33</sup> | N/A |
| Primary hMPCs were isolated from the gingiva of the individuals | Ren et al.<sup>67</sup> | N/A |
| HEK293T | ATCC | N/A |
| Human primary dermal fibroblasts | Zou et al.<sup>63</sup> | N/A |

| **Experimental Models: Organisms/Strains** |  |  |
| --- | --- | --- |
| C57BL/6J mice | SiPeiFu (Beijing) Biotechnology Co., Ltd. | N/A |
| **Oligonucleotides** |  |  |
| Primers for qRT-PCR, ChIP-qPCR, and vector constructions, as well as probes for DNA/RNA-FISH see Table S5 | This study | N/A |
| **Recombinant DNA** |  |  |
| pLVTHM | A gift from Didier Trono | Addgene plasmid #12247 |
| psPAX2 | A gift from Didier Trono | Addgene plasmid #12260 |
| pMD2.G | A gift from Didier Trono | Addgene plasmid #12259 |
| lentiSAM v2 | A gift from Didier Trono | Addgene plasmid #75112 |
| lentiMPH v2 | A gift from Didier Trono | Addgene plasmid #89308 |
| dCas9-KRAB | A gift from Didier Trono | N/A |
| HERVK RVLP construct | This study | N/A |
| **Software and Algorithms** |  |  |
| ImageJ (version 1.8.0) | Schneider et al.<sup>89</sup> | https://imagej.net/Welcome |
| Leica LAS AF Lite (version 2.6.0) | Leica | https://leica-las-af-lite.updatestar.com/ |
| GraphPadPrism 8.0 | GraphPad Software Inc. | https://www.graphpad.com/ |
| Image Lab 4.0.1 | Bio-Rad | https://www.bio-rad.com/zh-cn/product/imagelab-software?ID=KRE6P5E8Z |
| Phenochart™ 1.0 | Phenochart | https://www.akoyabio.com/support/software/phenochart-whole-slide-viewer/ |
| ImageScope (version 12.3.0.5056) | Leica | https://www.leicabiosystems.com/en-de/digital-pathology/manage/aperio-imagescope/ |
| IMARIS 9.8 | OXFORD INSTRUMENTs | https://imaris.oxinst.cn/newrelease |
| ZEN 3.1 | ZEISS | https://www.microshop.zeiss.com/en/us/system/zen+software/software+zen/zen+-+basic+software/410135-1002-310 |
| Analyze 10.0 | AnalyzeDirect | https://www.analyzedirect.com |
| TrimGalore (version 0.4.5) | Babraham Bioinformatics | https://github.com/FelixKrueger/TrimGalore |
| SAMtools (version 1.6) | Li et al.<sup>90</sup> | https://github.com/samtools/samtools |
| R (version 4.0.2) | N/A | https://www.r-project.org/ |
| pheatmap (version 1.0.12) | N/A | https://cran.r-project.org/web/packages/pheatmap/ |
| ggplot2 (version 3.3.2) | N/A | https://cran.r-project.org/web/packages/ggplot2/ |
| GSEA (version 4.1) | Subramanian et al.<sup>91</sup> | http://software.broadinstitute.org/gsea/index.jsp |
| Fastp (version 0.19.10) | Chen et al.<sup>92</sup> | https://github.com/OpenGene/fastp |
| hisat2 (version 2.0.4) | Kim et al.<sup>93</sup> | https://ccb.jhu.edu/software/hisat2/index.shtml |
| HTSeq (version 0.11.0) | Anders et al.<sup>94</sup> | https://htseq.readthedocs.io/en/master/ |
| featureCounts (version 2.0.0) | Liao et al.<sup>95</sup> | http://subread.sourceforge.net/featureCounts.html |
| STAR (version 2.7.1b and 2.7.9a) | Dobin et al.<sup>96</sup> | https://github.com/alexdobin/STAR |
| TEtranscripts (version 2.2.1) | Jin et al.<sup>97</sup> | https://github.com/mhammell-laboratory/TEtranscripts |
| DESeq2 (version 1.29.8 and 1.26.0) | Love et al.<sup>98</sup> | https://bioconductor.org/packages/release/bioc/html/DESeq2.html |
| bsmap (version 2.90) | Xi and Li<sup>99</sup> | https://hpc.nih.gov/apps/bsmap.html |
| ggpubr (version 0.4.0) | N/A | https://cran.microsoft.com/snapshot/2021-09-26/web/packages/ggpubr/index.html |
| IGV (version 2.8.12) | Thorvaldsdottir et al.<sup>100</sup> | https://igv.org |

### RESOURCE AVAILABILITY

#### Lead Contact

Further information and requests for resources and reagents should be directed to and will be fulfilled by the Lead Contact Dr. Guang-Hui Liu.

#### Materials Availability

This study did not generate new unique reagents.

#### Data and Code Availability

All datasets have been deposited in the Genome Sequence Archive in the National Genomics Data Center with accession numbers as indicated. The raw RNA-seq data for prematurely senescent and RS hMPCs: CRA003282. The raw WGBS data for prematurely senescent and RS hMPCs: HRA001144.^39^ The raw WGBS data for vehicle or 5-AZA treated hMPCs: HRA001879. The raw RNA-seq data for WT hMPCs (P6) after transduction with lentivirus containing sgRNA targeting HERVK or NTC using a CRISPRa system: CRA003283. The RNA-seq raw data for liver tissues of young and old cynomolgus monkeys: CRA002965.^66^ The RNA-seq raw data for lung tissues of young and old cynomolgus monkeys: CRA002810^88^. Any additional information required to reanalyze the data reported in this work is available from the Lead Contact upon request.

### EXPERIMENTAL MODELS AND SUBJECT DETAILS

#### Animals and human samples and ethics

C57BL/6J mice purchased from SiPeiFu (Beijing) Biotechnology Co., Ltd, were maintained in the animal care facilities of Institute of Biophysics and Institute of Zoology, Chinese Academy of Sciences, and fed *ad libitum* with standard laboratory chow and water in ventilated cages under a 12 h light/dark cycle. All mouse experiments followed the Principles for the Application Format for Ethical Approval for Research Involving Animals and were approved in advance by the Institutional Animal Care and Use Committee of the Institute of Zoology, Chinese Academy of Sciences. Anaesthetization of mice was conducted with isoflurane and euthanization was performed with CO2 followed by cervical dislocation. All cynomolgus monkey experiments were conducted under the Principles for the Ethical Treatment of Non-Human Primates and approved by the Animal Care and Ethics Committee of Institute of Zoology, Chinese Academy of Sciences. The young and old cynomolgus monkeys used for the experiments were raised at the Xieerxin Biology Resource with accreditation by a Laboratory Animal Care accredited facility in Beijing, in compliance with all local and federal laws governing animal research.^101^ The cynomolgus monkeys used in this study were the same ones reported in the previous studies.^101,102^ The lung, liver and skin samples were collected from 8 young (4-6 years old, 4 female and 4 male) and 8 old (18-21 years old, 4 female and 4 male) cynomolgus monkeys without clinical or experimental history. The HGPS cynomolgus monkeys were described in a previous study.^62^ Primary hMPCs were isolated from the gingiva of individuals of the indicated age under the approval from the Ethics Committee of the 306 Hospital of PLA in Beijing.^67^ Human eyelid skin samples were obtained from blepharoplasty of the indicated healthy donors under ethical approval of Peking Union Medical College Hospital Institutional Review Board.^63^ Human serum samples were collected under the approval of the Research Ethics Committee of the First Hospital of Kunming Medical University^88^. All human samples were obtained from healthy young and old donors. The age information of the human donors is listed in Table S2.

#### Cell culture

HEK293T cells and human fibroblasts were cultured in Dulbecco’s Modified Eagle Medium (DMEM, Thermo Fisher Scientific) supplemented with 10% fetal bovine serum (FBS, GIBCO, Thermo Fisher Scientific), 2 mM GlutaMAX (Thermo Fisher Scientific), 0.1 mM non-essential amino acids (NEAA, Thermo Fisher Scientific), 1% penicillin/streptomycin (Thermo Fisher Scientific). Human MPCs (hMPCs) were grown on 0.1% gelatin (Sigma Aldrich)-coated plates (CORNING) in hMPC culture medium containing 90% MEMα (Thermo Fisher Scientific), 10% FBS, 2 mM GlutaMAX, 0.1 mM NEAA, 1% penicillin/streptomycin and 1 ng/mL bFGF (Joint Protein Central). Cells were cultured in an incubator (Thermo Fisher Scientific) at 37°C with 5% CO_2_. WT hMPCs are replicatively senescent in their late passage (LP, P > 12), and prematurely senescent HGPS (*LMNA*^G608G/+^) and WS (*WRN*^-/-^) hMPC models exhibit growth arrest at passage 8-9 (referred to as LP in the premature aging model). HGPS and WS models were constructed via genome-editing in human embryonic stem cells (hESCs), followed by directed differentiation to hMPCs.^33^ Human fibroblasts are replicatively senescent in their late passage (LP, P > 23).

### METHOD DETAILS

#### Animal experiments

To evaluate whether lentiviral knockdown of MMTV alleviates aging-associated articular degeneration in mice, lentiviruses containing CRISPR-dCas9-KRAB-expressing empty backbone (CRISPRi-Control) or lentiviruses containing CRISPR-dCas9-KRAB and MMTV single-guide (sg) RNA (CRISRRi-sgMMTV) were injected into the articular cavities of 21-month-old mice twice with an interval of 4 weeks (n = 12 mice for each group). Grip strength was measured at 8 weeks after the first injection. To examine the effects of Abacavir on aging-associated articular degeneration, Abacavir was dissolved in PBS containing 1% DMSO at a final concentration of 5 mg/mL, and injected into the articular cavities of 22-month-old mice (n = 12 mice for each group) with a volume of 10 μL. Grip strength was measured at 8 weeks after the first injection. For long-term oral administration experiments, 18-month-old mice were treated daily with Abacavir dissolved in drinking water containing 0.4% DMSO at a final concentration of 1 mg/mL for a total of 3 mL, with 0.4% DMSO in drinking water as control (n = 23 mice for each group). The body weight, back, fur, skin, cataract and physical scoring, as well as the grip strength of mice were evaluated at 6, 12 and 24 weeks after treatment with Abacavir. The Y maze test was performed at 24 weeks after Abacavir treatment.

#### Grip strength test

The grip strength of the mice was assessed using a Grip Strength Meter (Panlab Grid Strength Meter, LE902). Briefly, four limbs of the mouse were placed on the top of the Grip Strength Meter and pulled along the direction of the grid at a constant rate until the Grip Strength Meter was released by the mouse. This process was repeated 10 times and the peak pull force at each time was recorded on a digital force transducer. The mean values from 10 consecutive trials were taken as the grip strength of each mouse. The number of mice (n) in each group is indicated in the figures; for example, n = 23 mice for vehicle or Abacavir in the drinking water as shown in Figure 7N; n = 12 mice injected with lentiviruses expressing control or sgMMTV using a CRISPRi system into the articular cavities as shown in Figure S7I; and n = 12 mice injected with vehicle or Abacavir into the articular cavities as shown in Figure S7L.

#### Y maze test

The Y maze test was performed using a Y-shaped arena consisting of three arms in identical lengths with 120°angles between each arm, a camera and a computer with tracking software. Briefly, the mouse was placed at the end of any arm of the Y maze, and allowed to explore the maze freely for 5 min. The camera system recorded the animal behavioural changes for 5 min. The total number of arm entries and the number of turns (alternating) reflecting that the mouse entered the three arms in a row were used to evaluate the short-term spatial memory. The number of mice (n) in each group is indicated in the figures; for example, n = 20 mice for vehicle and n = 22 for Abacavir in the drinking water as shown in Figure 7P.

#### *In vivo* bone histomorphometric analysis: Micro-Computed Tomography (Micro-CT)

The Micro-Computed Tomography (Micro-CT) scanning was conducted using a PE Quantum FX (PerkinElmer) on living mice. Briefly, the mice were anesthetized with isoflurane and scanned using the following settings. The proximal part of the tibias and the distal part of the femurs were scanned in a Micro-CT scanner with an isotropic voxel size of 18 μm. The X-ray tube voltage was set at 90 kV and the current was set at 160 μA, with an exposure time of 1 s. Quantification of the bone density was performed on the knee joint region of each mouse using Analyze 10.0 software. The number of mice (n) in each group is indicated in the figures; for example, n = 12 mice injected with lentiviruses expressing control or sgMMTV using a CRISPRi system into the articular cavities as shown in Figure 7G; and n = 10 mice injected with vehicle or n = 8 mice injected with Abacavir into the articular cavities as shown in Figure 7L.

#### Western blotting

Cells were lysed in 1 × SDS buffer (100 mM Tris-HCl (pH = 6.8), 10% glycerol, 2% SDS and 2% 2-mercaptoethanol) and boiled at 105°C for 10 min. Protein concentration was measured with the BCA Kit (Dingguochangsheng Biotech). Cell lysates were subjected to SDS-PAGE electrophoresis and electrotransferred onto PVDF membranes (Millipore). Subsequently, the membranes were incubated with primary antibodies and then HRP-conjugated secondary antibodies. For chemiluminescent detection, western blots were incubated with substrates (substrate A: 0.2 mM coumaric acid, 1.25 mM luminal, 0.1 M Tris-HCl (pH = 8.5); substrate B: 3% H_2_O_2_), in a ratio of 1,000 μL of substrate A to 3 μL substrate B, or SuperSignal™ West Femto Maximum Sensitivity Substrate (Thermo Fisher Scientific). Imaging was performed with the Image Lab software on a ChemiDoc XRS+ system (Bio-Rad Laboratories, Inc.). Quantification of protein band intensity was performed using ImageJ. GAPDH was used as a loading control. Three independent experiments were performed (n = 3) for each assay. The number of animal (n) in each group is indicated in the figures; for example, n = 8 young and aged cynomolgus monkeys as shown in Figure 6B; n = 5 young and aged mice as shown in Figure S7B. The antibodies used for western blotting are listed in the key resources table.

#### Immunoprecipitation (IP) of HERVK from conditioned medium

Cells were cultured in hMPC medium supplemented with 10% microvesicle-depleted fetal bovine serum (dFBS) for 48 h, which was prepared via ultracentrifugation at 100,000 × *g* for 16 h.^103^ Approximately 40 mL of conditioned medium from senescent cells was collected and filtered through a 0.2-μm filter (Pall Corporation) and pre-cleaned with Protein A/G Plus-Agarose (Santa Cruz Biotechnology) at 4°C for 2-3 h. Then, the harvested supernatants were incubated with a 1/1,000 dilution of HERVK-Env antibody (Austral Biologicals) and fresh Protein A/G Plus-Agarose at 4°C overnight. Finally, the supernatants were collected, and the beads were washed with PBS and subjected to western blotting.

#### Purification of microvesicle particles from conditioned medium

Cells were cultured in hMPC medium supplemented with 10% dFBS for 48 h before medium collection.^103^ The conditioned medium was collected and the corresponding cell number was counted. Medium from the same number of cells was used for microvesicle purification by centrifugation at 300 × *g* for 10 min to eliminate cell contamination and 15,000 × *g* for 20 min, followed by filtration through a 0.2-μm filter (Pall Corporation) and then ultracentrifugation at 110,000 × *g* for 2 h.^103^ Pellets were collected and resolved in an equal volume of PBS and subjected to viral RNA isolation or western blotting.

#### DNA/RNA isolation and quantitative (reverse transcriptase) PCR (q(RT-)PCR))

Total DNA from cells was extracted using the DNeasy Blood & Tissue Kit (Tiangen) following the manufacturer’s instructions and then processed for qPCR analysis. Total cellular RNA was extracted using TRIzol (Thermo Fisher Scientific), subjected to DNase I treatment to remove genomic DNA contamination, and reverse transcribed to cDNA using the GoScript Reverse Transcription System (Promega). q(RT-)PCR was performed using the THUNDERBIRD qPCR Mix (TOYOBO) on a CFX384 Real-Time System (Bio-Rad Laboratories, Inc.). 5S rDNA was used as an internal control for qPCR analysis of DNA, and *ACTB* or *GAPDH* was used as an internal control for qRT-PCR analysis of RNA. The number of animals or donors (n) in each group is indicated in the figures; for example, n = 4 human donors of young or old primary hMPCs as shown in Figure 2I; n = 8 young or aged cynomolgus monkeys as shown in Figures S6A, S6C ad S6D; n = 11 mice injected with lentiviruses expressing control or n = 12 mice injected with lentiviruses expressing sgMMTV using a CRISPRi system into the articular cavities as shown in Figure S7H; and n = 10 mice injected with vehicle, or n = 8 mice injected with Abacavir into the articular cavities as shown in Figure S7K. All primers used for qRT-PCR are listed in Table S5.

#### Viral RNA isolation from microvesicles

RNA was extracted from 140 μL of microvesicle suspension from the same number of cells using the QIAamp Viral RNA Mini Kit (QIAGEN) following the manufacturer’s instructions. Briefly, samples were incubated with pre-prepared buffer AVL and carrier RNA. After adding ethanol, the samples were loaded onto QIAamp Mini columns. After washing and elution, the eluted samples were treated with DNase I using a TURBO DNA-free Kit (Thermo Fisher Scientific). In brief, 2 μL DNase I and 2.4 μL reaction buffer were added to 20 μL eluted sample (viral RNA), followed by incubation at 37°C for 30 min. Subsequently, the DNase I was inactivated using the inactivation reagent and removed by centrifugation at 10,000 × *g* for 1.5 min. The collected supernatant was heated at 70°C for 10 min to inactive residual DNase I.

#### Droplet Digital PCR (ddPCR)

The purified viral RNA from microvesicles was reverse transcribed to cDNA using the GoScript Reverse Transcription System following the manufacturer’s instructions. Then, 20 μL of reaction solution containing cDNA templates, primers and QX200 ddPCR EvaGreen Supermix (Bio-Rad Laboratories, Inc.) were mixed with 70 μL QX200™ Droplet Generation Oil for EvaGreen (Bio-Rad Laboratories, Inc.) in a DG8 cartridge (Bio-Rad Laboratories, Inc.) using a QX200 droplet generator (Bio-Rad Laboratories, Inc.). The droplet was transferred to a 96-well plate and sealed with a PX1™ PCR Plate Sealer (Bio-Rad Laboratories, Inc.). After PCR amplification with a thermal cycler (Bio-Rad Laboratories, Inc.), the samples were subsequently subjected to a QX200 droplet reader (Bio-Rad Laboratories, Inc.) to analyze the absolute copy number of HERVK in microvesicles from the same number of cells in each group. ddH2O and cDNA from senescent hMPCs were used as negative and positive controls, respectively. Three biological replicates were performed for each group (n = 3).

### ELISA

For the human IL6 ELISA assay, the medium was collected and filtered with a 0.2-μm filter and incubated in an anti-IL6 antibody-coated plate (BioLegend) according to the manufacturer’s instructions. After incubation with a detection antibody, Avidin-HRP, freshly mixed TMB substrate, and stop solution, the plate was measured at 450 nm using a Synergy H1 Microplate Reader (BioTek). IL6 levels were normalized to the corresponding cell numbers. Three biological replicates were performed for each group (n = 3).

For HERVK-Env ELISA assay, conditioned medium or human serum was first concentrated into 100 or 10 times using Centricon Plus-70 (Millipore) or Amicon Ultra-15 (Millipore), respectively. The HERVK-Env protein in the conditioned medium or human serum was detected using the Human HERVK-Env Polyprotein ELISA Kit (CUSABIO) following the manufacturer’s instructions. Briefly, 100 μL concentrated medium or serum was added into the coated plate and incubated for 2 h at 37°C. After incubation with Biotin-antibody, Avidin-HRP, freshly mixed TMB substrate and stop solution, the plate was measured at 450 nm using a Synergy H1 Microplate Reader. HERVK-Env levels in the conditioned medium were normalized to the corresponding cell numbers. Three biological replicates were performed for each group (n = 3). HERVK-Env levels in human serum were also quantified, with n = 30 donors in each group.

A 2’3’-cGAMP ELISA Kit (Cayman) was used to measure the levels of 2’3’-cGAMP in cells. Briefly, 50 μL of cell lysates from the same number of indicated cell lines were loaded onto the plate followed by incubation with 2’3’-cGAMP HRP tracer and 2’3’-cGAMP polyclonal antiserum for 2 h at room temperature with shaking. After incubation with TMB substrate and stop solution, the plate was measured at 450 nm using a Synergy H1 Microplate Reader. 2’3’-cGAMP levels were normalized to the corresponding cell numbers. Three biological replicates were performed for each group (n = 3).

#### Immunofluorescence, immunohistochemistry staining and microscopy

For immunofluorescence staining, cells seeded on coverslips (Thermo Fisher Scientific) were washed twice with PBS, fixed in 4% paraformaldehyde (PFA), permeabilized in 0.4% Triton X-100 in PBS and blocked with 10% donkey serum. Coverslips were incubated with primary antibodies in blocking buffer at 4°C overnight and then with secondary antibodies for 1 h at room temperature. Nuclei were labeled with Hoechst 33342 (Thermo Fisher Scientific). The co-staining of HERVK-Env and ß-galactosidase was conducted using the Cellular Senescence Detection Kit-SPiDER-βGal (DOJINDO Laboratories). Briefly, cells seeded on coverslips were washed twice with PBS, fixed in 4% PFA for 3 min, and incubated with SPiDER-βGal for 30 min at 37°C. Then, the cells were washed with PBS and subjected to immunofluorescence staining of HERVK-Env following the normal procedure. Images were taken with a Leica SP5 confocal microscope or ZEISS confocal LSM900. The Z-stack 3D reconstruction of images was performed using IMARIS. For quantification of Ki67-positive cells, three biological replicates were set for each group (n = 3). Over 100 cells were quantified in each replicate. The quantification of HERVK-Env fluorescence intensity in each group was performed from more than 100 cells from 3 biological replicates.

Immunohistochemistry staining of tissue sections was performed using the DAB Staining Kit (ZSGB-BIO) according to the manufacturer’s instructions. Briefly, deparaffinized sections using Xylene and ethanol, were incubated with 3% H2O2 solution to block endogenous peroxidase activity before antigen retrieval using 10 mM sodium citrate buffer (pH = 6.0). Sections were then permeabilized, blocked and incubated with primary antibodies at 4°C overnight and secondary antibodies for 1 h, followed by staining with DAB substrate solution and Hematoxylin, dehydration and mounting. To remove the background, the sections were washed with PBS 5 times for 5 min each time. Images were taken with a LEICA Aperio CS2 LSM900. The white balance was adjusted before capturing the images. The ratios of positive cells were quantified. The number of animals or donors (n) in each group is indicated in the figures; for example, n = 5, 7 or 8 young or aged cynomolgus monkeys as shown in Figure 6C; n = 5, 10, or 20 fields from two sections of WT and HGPS cynomolgus monkeys as shown in Figure 6F; n = 5 or 6 young or aged human donors of skin as shown in Figure 6H; n = 8 young or aged mice as shown in Figures 7A; n = 5, 6 or 8 young or aged mice as shown in Figure S7C; n = 7 or 10 young or aged mice as shown in Figure S7D; n = 8 or 11 young or aged mice as shown in Figure S7E; n = 12 mice injected with lentiviruses expressing control or sgMMTV using a CRISPRi system into the articular cavities as shown in Figures 7C; n = 10 or 11 mice injected with lentiviruses expressing control or sgMMTV using a CRISPRi system into the articular cavities as shown in Figure 7D; n = 11 mice injected with lentiviruses expressing control or sgMMTV using a CRISPRi system into the articular cavities as shown in Figure 7E; n = 11 or 12 mice injected with lentiviruses expressing control or sgMMTV using a CRISPRi system into the articular cavities as shown in Figure S7G; and n = 8 or 9 mice injected with vehicle or Abacavir into the articular cavities as shown in Figures 7I; n = 7 or 8 mice injected with vehicle or Abacavir into the articular cavities as shown in Figure 7J; n = 8 or 10 mice injected with vehicle or Abacavir into the articular cavities as shown in Figure S7J.

Safranin-O/Fast Green staining was performed by Servicebio Technology Co., Ltd. (Beijing, China). Briefly, sections were baked at 60°C overnight and cooled at room temperature for 20-30 min followed by deparaffinization and hydration in 70% ethanol. Then, the slides were stained with Weigert’s Iron Hematoxylin for 7 min, 0.08% Fast Green for 3 min, and rinsed in 1.0% acetic acid for 10 s. After staining with 0.1% Safranin-O for 5 min and rinsing in 0.5% acetic acid and distilled water, sections were mounted. Images were taken with a LEICA Aperio CS2 LSM900. The mean values from at least 6 regions of articular cartilages were taken as the thickness of articular cartilages of each mouse. The number of mice in each group is indicated in the figures; for example, n = 8 young or n = 11 aged mice as shown in Figure S7E; n = 12 mice injected with lentiviruses expressing control or sgMMTV using a CRISPRi system into the articular cavities as shown in Figure 7F; and n = 10 mice injected with vehicle or n = 8 mice injected with Abacavir into the articular cavities as shown in Figure 7K.

Antibodies used for immunofluorescence and immunohistochemistry staining are listed in the key resources table.

#### (sm)RNA/DNA-FISH

RNA-FISH was performed using the QuantiGene^®^ ViewRNA ISH Cell Assay Kit (Thermo Fisher Scientific) following the manufacturer’s instructions. Briefly, cells seeded on coverslips (Thermo Fisher Scientific) were fixed in fresh 4% formaldehyde solution, permeabilized with Detergent Solution QC and digested with Protease QC. Probe HERVK Alexa Fluor 488 was diluted in Probe Set Diluent QF and added to wells for 3 h at 41 + 1°C in an HB-1000 Hybridizer (Gene Company Limited). Coverslips were incubated with Working Pre-Amplifier Mix Solution for 30 min, Working Amplifier Mix Solution for 30 min and Working Label Probe Mix Solution for 30 min at 40 + 1°C in an HB-1000 Hybridizer, followed by PBS washing in each step. Nuclei were labeled with DAPI and slides were mounted and subjected to imaging with a Leica SP5 confocal microscope. HERVK RNA-FISH fluorescence intensity was quantified from more than 150 cells from 3 biological replicates.

Detection of HERVK mRNA and DNA in cytoplasm was performed via single-molecule (sm)-FISH. The targeting sequences of the probes for HERVK were split into pairs to increase the detection specificity, and probe pairs against HERVK were designed by Spatial FISH, Co., Ltd. Briefly, cells were fixed in 4% PFA in DEPC-treated PBS at room temperature for 20 min and washed for three times with DEPC-treated PBS, followed by 70% ethanol immersion for at least 24 h. Then, cells were treated with 0.2 M HCl in DEPC-treated H2O for 10 min and 5 μg/mL protein kinase K in DEPC-treated PBS for 5 min, respectively. For mRNA detection, cells were directly incubated with targeting probe pairs (10 μM for each probe) in hybridization buffer (10% deionized formamide, 2 × SSC, 10% dextran sulfate, and 2 mM VRC) at 42°C overnight. For detection of HERVK DNA, the cells were sequentially treated with RNase A (100 μg/mL in 2 × SSC) at 37°C for 1 h, 70% formamide in 2 × SSC at 72°C for 10 min, and a pre-cooled ethanol gradient (80%, 90%, 100%) for 1 min each gradient, followed by incubation with targeting probe pairs (10 μM for each probe) in hybridization buffer at 42°C overnight. After washing with washing buffer (2 × SSC, 30% formamide, 0.1% Triton-X 100, 2 mM VRC) three times, singlemolecule detection of HERVK mRNA and DNA was realized using the following probes including amplification probes (SFSP-001 and SFTP-001 from Spatial FISH, Co., Ltd) and fluorescent probe (SFFP-001, Spatial FISH, Co., Ltd). Finally, the cells were counterstained with DAPI and imaged with a Leica SP5 confocal microscope or N-STORM. The relative number of fluorescent spots of HERVK DNA in the cytoplasm was quantified from over 10 cells in each sample. The targeting sequences of the probe pairs are listed in Table S5.

#### (Immuno-)Transmission electron microscopy (TEM)

Cells were pelleted and fixed with 2.5% (vol/vol) glutaraldehyde with Phosphate Buffer (PB) (0.1 M, pH = 7.4) at room temperature for 20 min and then at 4°C overnight. Routine heavy metal staining was conducted. Briefly, cells were postfixed with 1% (wt/vol) osmium tetraoxide in PB at 4°C for 2 h, dehydrated through a graded ethanol series into pure acetone, infiltrated in a graded mixture of acetone and resin, and then embedded in pure resin with 1.5% BDMA and polymerized at 45°C for 12 h, and then at 60°C for 48 h. Ultrathin sections (70-nm thick) were sectioned with a microtome (Leica EM UC6), and double-stained with uranyl acetate and lead citrate. For Immuno-TEM, cells seeded in 35-mm petri dishes (CORNING) were prefixed with 2% formaldehyde in warmed culture medium at 37°C in a cell incubator for 10 min, and with 4% formaldehyde at room temperature for 1 h and 4°C overnight, and then transferred into 1% formaldehyde. Samples were dehydrated through an ethanol gradient, infiltrated with a LR Gold resin gradient, and embedded in LR Gold resin containing initiator. Samples were then polymerized by UV light at −20°C for 24 h using a Leica AFS2 and sectioned using a Leica

EM UC7. For staining, sections were blocked with 2% BSA in PB (Jackson ImmunoResearch), incubated with anti-HERVK-Env antibody at room temperature for 2 h, washed with PB buffer containing 0.1% cold fresh gelatin for 2 min for five times, incubated with 6-nm gold-labeled anti-mouse secondary antibody (Jackson ImmunoResearch) at room temperature for 1 h, and then fixed with 1% glutaraldehyde in ddH2O. After washing with ddH2O, sections were stained with 2% uranyl acetate at room temperature for 30 min and then processed for imaging using a TEM Spirit 120 kV (FEI Tecnai Spirit 120 kV). The number of RVLPs or HERVK RVLPs labeled with gold particles per cell was quantified from more than 40 cells in each group.

#### Plasmid construction

To generate HERVK^44,45^ or STING knockdown vectors,^8^ specific shRNAs were cloned into MluI/ClaI sites of the lentiviral vector pLVTHM (Addgene, #12247). To activate endogenous HERVK, nontargeting control sgRNA (sgNTC)^104^ or sgRNA targeting HERVK LTR (sgHERVK)^42^ was cloned into lentiSAM v2 vector (Addgene, #75112) via the ESP3I site, and then co-transfected into hMPCs with lentiMPH v2 (Addgene, #89308). The constructs that repress endogenous HERVK using a dCas9-KRAB-based system (CRISPRi-sgHERVK) and dCas9-KRAB-expressing empty backbone (CRISPRi-control) were kind gifts from the Dr. Didier Trono’s lab (Ecole Polytechnique Fédérale de Lausanne (EPFL), Lausanne, Switzerland).^43^ We investigated the target specificity of sgRNAs used for HERVK CRISPRa and CRISPRi systems. We first obtained the genomic locations of RepeatMasker-annotated repetitive elements based on the hg19 assembly, and then constructed the repetitive element BLAST (Basic Local Alignment Search Tool) database for *LTR5_Hs* (the promoter of HERVK). By comparing the sgRNA sequence with the constructed *LTR5_Hs* database, we found that ~30% of total *LTR5_Hs* elements showed good matches (expected value [E-value] < 1 × 10^-5^) and were potentially targeted by CRISPRa and CRISPRi sgRNAs, which is highly consistent with the targeting efficiency of sgRNAs designed for HERVK transcriptional activation and/or repression in recently published studies.^105,106^ To repress endogenous MMTV, the sgRNAs targeting the LTR region of MMTV (GenBank: AB049193.1) were designed by http://chopchop.cbu.uib.no/, and cloned into a dCas9-KRAB-expressing empty backbone via the BsmB1 site (CRISPRi-sgMMTV). To investigate the specificity of the sgRNA for MMTV, we first obtained the genomic locations for endogenous MMTVs annotated by RepeatMasker based on the mouse assembly (mm10 version) and found three potentially full-length MMTVs. By comparing the sgRNA sequence with the mouse LTR/ERVK elements, we found that the designed CRISPRi sgRNA showed high potential to target two of three potential full-length MMTVs (E-value < 0.05). The targeting sequences of shRNA or sgRNA are listed in Table S5.

To construct a plasmid producing HERVK RVLPs, full-length HERVK was synthesized according to a previously published sequence with NheI and ApaI sites at the N- and C-termini, respectively.^57,107^ The KpnI site of pEGFP-N3 was disrupted using a Fast Mutagenesis System (TransGen), termed pEGFP-N3-KpnI’. HERVK was cloned into pEGFP-N3-KpnI’ via NheI/ApaI sites, and its expression was driven by the CMV promotor of the vector. Of note, due to the stop signal in the 3’-LTR of HERVK, GFP in the vector backbone is not expressed in pricinple. Another CMV-EGFP cassette was PCR amplified from pEGFP-C1 and inserted into the KpnI site of pEGFP-N3-HERVK within the Env region of HERVK, which was designed following previously published studies.^58,108^ To boost HERVK virus protein expression and improve RVLP production,^58,108^ other HERVK expression plasmids including gag-pro-pol, env or rev fragments, were generated by PCR amplification using synthesized HERVK as the template, which was cloned into the pEGFP-N3 vector via NheI/ApaI sites for the Env fragment or KpnI/BamHI sites for the gag-pro-pol and rev fragments. Of note, to amplify the rev fragment, the reverse primer was designed to cover the additional 63 nucleotides at the C-terminal. In principle, the full-length HERVK containing LTR with the new CMV-EGFP cassette within the Env region could be packaged into RVLPs, but not other fragments.

Sequences of the sgRNAs used for CRISPRa and CRISPRi, as well as primers used for plasmid construction are listed in Table S5.

#### Production of HERVK RVLPs and lentiviruses

The production and purification of HERVK RVLPs were conducted by transfecting HEK293T cells with the full-length HERVK construct using Lipofectamine 3000 Transfection Reagent (Thermo Fisher Scientific). To boost the viral protein expression of HERVK and improve the production and transfection capacity of RVLPs, full-length HERVK with the CMV-EGFP cassette was co-transfected with other HERVK expression plasmids (encoding gag-pro-pol, env and rev) as well as VSG plasmid into HEK293T cells. The supernatants containing HERVK RVLPs were harvested at 48 h and 72 h after transfection, filtered with a 0.2-μm filter, and then concentrated by ultracentrifugation at 100,000 ×*g* for 2 h. The vehicle was purified using the same procedure from the medium of HEK293T cells transfected with empty vector.

The lentivirus packaging was conducted according to previous studies.^70^ HEK293T cells were transfected with lentiviral vectors together with the packaging plasmids pMD2.G (Addgene, #12260) and psPAX2 (Addgene, #12259) using Lipofectamine 3000 Transfection Reagent. Supernatants containing lentiviruses were harvested at 48 h and 72 h after transfection, filtered with a 0.2-μm filter, and concentrated by ultracentrifugation at 19,400 rpm for 2.5 h. The virus pellets were suspended and assessed for viral titers. The purification of viruses was performed under the safety rules of the Biological Safety Levels (BSL)-2 laboratory.

#### Cell treatment

5-AZA treatment was performed according to previous publications.^46,47^ Briefly, WT hMPCs (P6) were seeded (recorded as passage 0 (P0) post-treatment) and treated with 500 ng/μL 5-AZA (dissolved in ddH2O) for one passage (around 4-6 days). The medium supplemented with 5-AZA was changed every two days. Then, the cells were passaged and cultured in fresh medium. At P2 post-treatment (6 days after 5-AZA treatment), cells were harvested for Ki67 immunofluorescence staining, SA-β-gal activity analysis and RNA/DNA extraction.

The Abacavir treatment assay was conducted as described in a previous study.^53^ Briefly, HGPS hMPCs (P7) were treated with different concentrations (0.2 μM, 1 μM and 10 μM) of Abacavir for two passages. During the treatment, cells were cultured with the culture medium being changed every two days, and the medium was supplemented with different concentrations (0.2 μM, 1 μM and 10 μM) of Abacavir. From the cell density and morphology, we found that treatment with the lowest concentration (0.2 μM) of Abacavir showed the best effect to alleviate premature senescence in HGPS hMPCs. Therefore, we used 0.2 μM as the final concentration for Abacavir treatment for the following experiments. After treatment, cells were collected for Ki67 immunofluorescence staining, SA-β-gal activity analysis and RNA/DNA extraction.

For infection with HERVK RVLPs, WT hMPCs (P6) were seeded (P0 post-treatment) and incubated with HERVK RVLPs in the presence of polybrene (Sigma-Aldrich) for 24 h. To increase the infection sensitivity of HERVK RVLPs, hMPCs were then subjected to centrifugation at 1,200 ×*g* for 2.5 h. After culture for two passages, cells were processed for Ki67 immunofluorescence staining, SA-β-gal activity analysis and RNA/DNA extraction. To block the HERVK RVLPs, pretreatment with IgG or anti-HERVK-Env for 30 min in a 37°C incubator was performed.

For lentivirus transduction, WT hMPCs (P6), HGPS or WS hMPCs (P5) were seeded (P0 post treatment) and incubated with lentiviruses in the presence of polybrene for 24 h. After two passages, cells were processed for Ki67 immunofluorescence staining, SA-β-gal activity analysis and RNA/DNA extraction.

For conditioned medium treatment, 50% medium collected from the indicated cell culture with 50% fresh medium was mixed and used for culturing early-passage WT hMPCs (P5) in the presence of polybrene (Sigma-Aldrich) with the medium being changed every two days. The SC-CM (HGPS (P7), WS (P7) and WT (LP, P14)) also accelerated young hMPC senescence without polybrene, but it took a longer time to show effects. The cells were passaged and then collected for Ki67 immunofluorescence staining, SA-β-gal activity analysis and RNA/DNA extraction. To detect the RVLPs from conditioned medium adhering to the surface of young cells, cells were treated with conditioned medium for 12 h and subjected to TEM analysis.

To culture the cells with pooled human serum from young (18-25 years old, n = 5) or old individuals (65-80 years old, n = 5), primary hMPCs isolated from the gingiva of young donors were precultured in normal hMPC culture medium for 24 h. After washing with PBS three times, hMPCs were cultured in the medium containing 10% human serum for two passages and then collected for further analysis.

#### Clonal expansion assay

Five thousand cells were seeded in one well of a 6-well plate (Corning) precoated with gelatin (Sigma) and cultured for approximately 10 days. The cells were then fixed with 4% PFA for 30 min and stained with 10% crystal violet for 30 min. The relative cell integral density was calculated using ImageJ software. Three biological replicates are performed for each group (n = 3).

#### SA-β-gal staining

SA-β-gal staining was performed as described previously.^33,109^ In brief, cells were fixed with a buffer containing 2% (w/v) formaldehyde and 0.2% (w/v) glutaraldehyde for 5 min and incubated with staining buffer containing 1 mg/mL X-gal at 37°C overnight. The optical microscope was used to observe SA-β-gal-stained cells and the percentage of SA-β-gal-positive cells was calculated using ImageJ software. Three biological replicates are performed for each group (n = 3). Over 100 cells were quantified in each replicate.

#### ChIP-qPCR

ChIP-qPCR was performed as previously described^110^ with some minor modifications. Briefly, cells were fixed in 1% formaldehyde in PBS for 10 min at room temperature and then quenched by 125 mM glycine. Then, cells were lysed on ice for 10 min and subjected to sonication using a Covaris S220 Focused-ultrasonicator. The collected supernatants were incubated with Dynabeads Protein A (Thermo Fisher Scientific) pre-conjugated with 2.3 μg indicated antibodies or IgG at 4°C overnight. After washing, samples were digested with proteinase K (New England Biolabs) and reverse crosslinked at 68°C for 2 h on a thermomixer. DNA was extracted using phenol-chloroform-isoamyl alcohol and subjected to qPCR analysis. Three biological replicates were performed for each group (n = 3). Four technical replicates were performed for each biological replicate. For immunoprecipitation of cGAS, the cytoplasmic fraction of cells extracted after fixation^11^ was incubated with anti-cGAS antibody- or IgG-conjugated Dynabeads Protein A at 4°C overnight. The DNA extraction procedure was performed as above. qPCR analysis of 5S rDNA, which was in principle absent in the cytoplasmic fraction, was used to exclude the nuclear genomic contamination. Four technical replicates were performed for each group (n = 4). The ChIP-qPCR data are presented as fold enrichment. Briefly, the negative control (IgG) sample is given a value of ‘1’, and everything else will then be normalized as a fold change of this negative control sample. The primers used for ChIP-qPCR are listed in Table S5.

#### Strand-specific RNA sequencing (RNA-seq)

Strand-specific RNA-seq libraries were prepared and sequenced by Novogene Bioinformatics Technology Co. Ltd. In brief, total RNA was extracted from 1 × 10^6^ cells per duplicate using TRIzol reagent, and genomic DNA was removed. RNA concentrations were measured using a Qubit^™^ RNA Assay Kit with a Qubit^®^2.0 Fluorometer (Thermo Fisher Scientific). A total amount of 3 μg RNA per sample was used for the RNA sample preparations and library constructions. Then, ribosomal RNA was removed using the Epicentre Ribo-Zero™rRNA Removal Kit (Epicentre), and rRNA-free residue was discarded by ethanol precipitation. Sequencing libraries were prepared using the NEBNext^®^Ultra™ Directional RNA Library Prep Kit for Illumina^®^(New England Biolabs) following the manufacturer’s instructions. High-throughput sequencing was performed on an Illumina NovaSeq 6000 platform.

#### RNA-seq data processing

The processing pipeline for RNA-seq data has been reported previously.^71,101^ Pair-end raw reads were trimmed by TrimGalore (version 0.4.5) (Babraham Bioinformatics) (https://github.com/FelixKrueger/TrimGalore) and mapped to the human (*Homo sapiens*) hg19 or cynomolgus macaque *(Macaca Fascicularis)* MacFas5.0 reference genome obtained from the UCSC genome browser database using STAR (version 2.7.1a) or hisat2 (version 2.0.4).^93,96^ High-quality mapped reads (score of mapping quality more than 20) were then used for counting reads by HTSeq (version 0.11.0) or featureCounts (version 1.6.4).^94,95^ Differentially expressed genes (DEGs) were calculated by the DESeq2 R package (version 1.29.8) with the cutoff “|log_2_(fold change)| > 0.5 and adjusted p value < 0.05” for hMPCs.^98^ SASP genes were obtained from a previous study.^9^ Gene set enrichment analysis (GSEA) was conducted by GSEA application (version 2.2.4).^91^

#### Analysis of the expression levels of repetitive elements

To evaluate the expression levels of repetitive elements, the cleaned reads were mapped to the human *(Homo sapiens)* hg19 reference genome using STAR software (version 2.7.9a) with the parameter “--outFilterType BySJout --winAnchorMultimapNmax 100 --outFilterMultimapNmax 100 --outFilterMismatchNoverLmax 0.04”.^96^ Transposable element quantification and the differential analysis were computed using the TEtranscripts software (version 2.2.1).^97^ In brief, TEtranscripts simultaneously counted the gene abundances and transposon abundances and utilized R package DESeq2 (version 1.26.0) for the differential analysis. Then, differentially expressed repetitive elements were computed with a cutoff “|log2(fold change)| > 0.2 and FDR < 0.05”. Class for repetitive elements was annotated within RepeatMasker, which can be classified as LTR, LINE, SINE, DNA (also known as DNA transposons), satellite, and some RNA repeats.

#### Whole genome bisulfite sequencing (WGBS) library construction and sequencing

Library preparation and sequencing for WGBS were performed as previously reported^111^. In brief, genomic DNA was extracted from 2 × 10^6^ cells per duplicate using DNeasy Blood & Tissue Kits (QIAGEN) and sheared to 100-300 bp with a sonicator. Then, bisulfite treatment, library preparation, quality control and sequencing were conducted by Novogene Bioinformatics Technology Co. Ltd.

#### WGBS data processing

WGBS data analysis was performed as previously reported^111^. In brief, raw sequencing reads were trimmed by fastp software (version 0.19.10) with default parameters.^92^ Then, cleaned reads were mapped to the human *(Homo sapiens)* hg19 reference genome obtained from the UCSC genome browser database using bsmap (version 2.90) with parameters “-v 0.1 -g 1 -R -u”. CpG DNA methylation levels for each cytosine site were calculated by the methratio program provided by bsmap. To ensure the accuracy of methylation level detection, forward and reverse strand reads for each CpG site were combined and only CpG sites with a depth of more than 5 were kept for downstream analysis. To calculate the relative CpG DNA methylation level in repetitive element loci, we obtained the genomic loci for each RepeatMasker-annotated repetitive element from the UCSC genome browser. Then, the average CpG DNA methylation level for each RepeatMasker-annotated repetitive element was calculated, and the statistical analysis was conducted by the ggpubr R package (version 0.4.0) in R (version 4.0.2).

#### Quantification and statistical analysis

All data were statistically analyzed using the PRISM version 8 software (GraphPad Software). Results are presented as the mean ± SEM. Comparisons were conducted using the two-tailed student’s t test or one-way ANOVA. p values < 0.05 were considered statistically significant (*), p values < 0.01 and p values < 0.001 were considered highly statistically significant (** and ***).

## Supplementary Table Legends

**Table S1.** Differentially expressed repetitive elements in replicatively and prematurely senescent hMPCs, related to Figure 1.

**Table S2.** Sample information used in this study, related to Figures 2 and 6.

**Table S3.** Differentially expressed genes in replicatively and prematurely senescent hMPCs, related to Figure 4.

**Table S4.** Differentially expressed genes in WT hMPCs (P6) transduced with lentiviruses expressing non-targeting control sgRNA (sgNTC) or sgRNA targeting HERVK (sgHERVK) using a CRISPR activation system (CRISPRa), related to Figure 4.

**Table S5.** List of primers and probes used in this study, related to STAR Methods.

## REFERENCES

1. Zhang, W., Qu, J., Liu, G.H., and Belmonte, J.C.I. The ageing epigenome and its rejuvenation. Nature reviews Molecular cell biology 2020; 21:137–150

2. Lopez-Otin, C., and Kroemer, G. Hallmarks of Health. Cell 2020; 10.1016/j.cell.2020.11.034

3. Horvath, S., and Raj, K. DNA methylation-based biomarkers and the epigenetic clock theory of ageing. Nature reviews Genetics 2018; 19:371–384

4. Kennedy, B.K., Berger, S.L., Brunet, A., Campisi, J., Cuervo, A.M., Epel, E.S., Franceschi, C., Lithgow, G.J., Morimoto, R.I., Pessin, J.E., et al. Geroscience: linking aging to chronic disease. Cell 2014; 159:709–713

5. Sun, Y., Li, Q., and Kirkland, J.L. Targeting senescent cells for a healthier longevity: the roadmap for an era of global aging. Life Medicine 2022; 10.1093/lifemedi/lnac030

6. Kang Wang, H.L., Qinchao Hu1, Lingna Wang, Jiaqing Liu, Zikai Zheng, Weiqi Zhang, Jie Ren, Fangfang Zhu and Guang-Hui Liu Epigenetic regulation of aging: implications for interventions of aging and diseases. Signal Transduction and Targeted Therapy 2022; 10.1038/s41392-022-01211-8

7. Cai, Y., Song, W., Li, J., Jing, Y., Liang, C., Zhang, L., Zhang, X., Zhang, W., Liu, B., An, Y., et al. The landscape of aging. Science China Life sciences 2022; 10.1007/s11427-022-2161-3

8. Bi, S., Liu, Z., Wu, Z., Wang, Z., Liu, X., Wang, S., Ren, J., Yao, Y., Zhang, W., Song, M., et al. SIRT7 antagonizes human stem cell aging as a heterochromatin stabilizer. Protein & cell 2020; 10.1007/s13238-020-00728-4

9. De Cecco, M., Ito, T., Petrashen, A.P., Elias, A.E., Skvir, N.J., Criscione, S.W., Caligiana, A., Brocculi, G., Adney, E.M., Boeke, J.D., et al. L1 drives IFN in senescent cells and promotes age-associated inflammation. Nature 2019; 566:73–78

10. Liang, C., Ke, Q., Liu, Z., Ren, J., Zhang, W., Hu, J., Wang, Z., Chen, H., Xia, K., Lai, X., et al. BMAL1 moonlighting as a gatekeeper for LINE1 repression and cellular senescence in primates. Nucleic acids research 2022; 50:3323–3347

11. Simon, M., Van Meter, M., Ablaeva, J., Ke, Z., Gonzalez, R.S., Taguchi, T., De Cecco, M., Leonova, K.I., Kogan, V., Helfand, S.L., et al. LINE1 Derepression in Aged Wild-Type and SIRT6-Deficient Mice Drives Inflammation. Cell metabolism 2019; 29:871–885.e875

12. Van Meter, M., Kashyap, M., Rezazadeh, S., Geneva, A.J., Morello, T.D., Seluanov, A., and Gorbunova, V. SIRT6 represses LINE1 retrotransposons by ribosylating KAP1 but this repression fails with stress and age. Nature communications 2014; 5:5011

13. Dubnau, J. The Retrotransposon storm and the dangers of a Collyer’s genome. Current opinion in genetics & development 2018; 49:95–105

14. Gorbunova, V., Seluanov, A., Mita, P., McKerrow, W., Fenyö, D., Boeke, J.D., Linker, S.B., Gage, F.H., Kreiling, J.A., Petrashen, A.P., et al. The role of retrotransposable elements in ageing and age-associated diseases. Nature 2021; 596:43–53

15. Liu, B., Qu, J., Zhang, W., Izpisua Belmonte, J.C., and Liu, G.H. A stem cell aging framework, from mechanisms to interventions. Cell reports 2022; 41:111451

16. Johnson, W.E. Origins and evolutionary consequences of ancient endogenous retroviruses. Nature reviews Microbiology 2019; 17:355–370

17. Blikstad, V., Benachenhou, F., Sperber, G.O., and Blomberg, J. Evolution of human endogenous retroviral sequences: a conceptual account. Cellular and molecular life sciences: CMLS 2008; 65:3348–3365

18. Subramanian, R.P., Wildschutte, J.H., Russo, C., and Coffin, J.M. Identification, characterization, and comparative genomic distribution of the HERV-K (HML-2) group of human endogenous retroviruses. Retrovirology 2011; 8:90

19. Marchi, E., Kanapin, A., Magiorkinis, G., and Belshaw, R. Unfixed endogenous retroviral insertions in the human population. Journal of virology 2014; 88:9529–9537

20. Stoye, J.P. Studies of endogenous retroviruses reveal a continuing evolutionary saga. Nature reviews Microbiology 2012; 10:395–406

21. Garcia-Montojo, M., Doucet-O’Hare, T., Henderson, L., and Nath, A. Human endogenous retrovirus-K (HML-2): a comprehensive review. Critical reviews in microbiology 2018; 44:715–738

22. Vargiu, L., Rodriguez-Tome, P., Sperber, G.O., Cadeddu, M., Grandi, N., Blikstad, V., Tramontano, E., and Blomberg, J. Classification and characterization of human endogenous retroviruses; mosaic forms are common. Retrovirology 2016; 13:7

23. Dendrou, C.A., Fugger, L., and Friese, M.A. Immunopathology of multiple sclerosis. Nature reviews Immunology 2015; 15:545–558

24. Sankowski, R., Strohl, J.J., Huerta, T.S., Nasiri, E., Mazzarello, A.N., D’Abramo, C., Cheng, K.F., Staszewski, O., Prinz, M., Huerta, P.T., et al. Endogenous retroviruses are associated with hippocampus-based memory impairment. Proceedings of the National Academy of Sciences of the United States of America 2019; 116:25982–25990

25. Mameli, G., Erre, G.L., Caggiu, E., Mura, S., Cossu, D., Bo, M., Cadoni, M.L., Piras, A., Mundula, N., Colombo, E., et al. Identification of a HERV-K env surface peptide highly recognized in Rheumatoid Arthritis (RA) patients: a cross-sectional case-control study. Clinical and experimental immunology 2017; 189:127–131

26. Bieda, K., Hoffmann, A., and Boller, K. Phenotypic heterogeneity of human endogenous retrovirus particles produced by teratocarcinoma cell lines. The Journal of general virology 2001; 82:591–596

27. Qu, Y., Izsvák, Z., and Wang, J. Retrotransposon: A Versatile Player in Human Preimplantation Development and Health. Life Medicine 2022; 10.1093/lifemedi/lnac041

28. Hurst, T.P., and Magiorkinis, G. Epigenetic Control of Human Endogenous Retrovirus Expression: Focus on Regulation of Long-Terminal Repeats (LTRs). Viruses 2017; 9:

29. Tie, C.H., and Rowe, H.M. Epigenetic control of retrotransposons in adult tissues: implications for immune regulation. Current opinion in virology 2017; 25:28–33

30. Kudlow, B.A., Kennedy, B.K., and Monnat, R.J., Jr. Werner and Hutchinson-Gilford progeria syndromes: mechanistic basis of human progeroid diseases. Nature reviews Molecular cell biology 2007; 8:394–404

31. Liu, G.H., Barkho, B.Z., Ruiz, S., Diep, D., Qu, J., Yang, S.L., Panopoulos, A.D., Suzuki, K., Kurian, L., Walsh, C, et al. Recapitulation of premature ageing with iPSCs from Hutchinson-Gilford progeria syndrome. Nature 2011; 472:221–225

32. Zhang, W., Li, J., Suzuki, K., Qu, J., Wang, P., Zhou, J., Liu, X., Ren, R., Xu, X., Ocampo, A., et al. Aging stem cells. A Werner syndrome stem cell model unveils heterochromatin alterations as a driver of human aging. Science (New York, NY) 2015; 348:1160–1163

33. Wu, Z., Zhang, W., Song, M., Wang, W., Wei, G., Li, W., Lei, J., Huang, Y., Sang, Y., Chan, P., et al. Differential stem cell aging kinetics in Hutchinson-Gilford progeria syndrome and Werner syndrome. Protein & cell 2018; 9:333–350

34. Geng, L., Liu, Z., Zhang, W., Li, W., Wu, Z., Wang, W., Ren, R., Su, Y., Wang, P., Sun, L., et al. Chemical screen identifies a geroprotective role of quercetin in premature aging. Protein & cell 2019; 10:417–435

35. Li, Y., Zhang, W., Chang, L., Han, Y., Sun, L., Gong, X., Tang, H., Liu, Z., Deng, H., Ye, Y., et al. Vitamin C alleviates aging defects in a stem cell model for Werner syndrome. Protein & cell 2016; 7:478–488

36. Campisi, J., Kapahi, P., Lithgow, G.J., Melov, S., Newman, J.C., and Verdin, E. From discoveries in ageing research to therapeutics for healthy ageing. Nature 2019; 571:183–192

37. W. Wang, Y.Z., S. Sun, W. Li, M. Song, Q. Ji, Z. Wu, Z. Liu, Y. Fan, F. Liu, J. Li, C. R. Esteban, S. Wang, Q. Zhou, J. C. I. Belmonte, W. Zhang, J. Qu, F. Tang, G.-H. Liu. A genomewide CRISPR-based screen identifies KAT7 as a driver of cellular senescence. Transl Med 2021; 13:

38. Robin, J.D., and Magdinier, F. Physiological and Pathological Aging Affects Chromatin Dynamics, Structure and Function at the Nuclear Edge. Frontiers in genetics 2016; 7:153

39. Liu, Z., Ji, Q., Ren, J., Yan, P., Wu, Z., Wang, S., Sun, L., Wang, Z., Li, J., Sun, G., et al. Large-scale chromatin reorganization reactivates placenta-specific genes that drive cellular aging. Developmental cell 2022; 57:1347-1368.e1312

40. Zhao, D., and Chen, S. Failures at every level: breakdown of the epigenetic machinery of aging. Life Medicine 2022; 10.1093/lifemedi/lnac016

41. Contreras-Galindo, R., Kaplan, M.H., Leissner, P., Verjat, T., Ferlenghi, I., Bagnoli, F., Giusti, F., Dosik, M.H., Hayes, D.F., Gitlin, S.D., et al. Human endogenous retrovirus K (HML-2) elements in the plasma of people with lymphoma and breast cancer. Journal of virology 2008; 82:9329–9336

42. Li, W., Lee, M.H., Henderson, L., Tyagi, R., Bachani, M., Steiner, J., Campanac, E., Hoffman, D.A., von Geldern, G., Johnson, K., et al. Human endogenous retrovirus-K contributes to motor neuron disease. Science translational medicine 2015; 7:307ra153

43. Turelli, P., Playfoot, C., Grun, D., Raclot, C., Pontis, J., Coudray, A., Thorball, C., Duc, J., Pankevich, E.V., Deplancke, B., et al. Primate-restricted KRAB zinc finger proteins and target retrotransposons control gene expression in human neurons. Science advances 2020; 6:eaba3200

44. Grow, E.J., Flynn, R.A., Chavez, S.L., Bayless, N.L., Wossidlo, M., Wesche, D.J., Martin, L., Ware, C.B., Blish, C.A., Chang, H.Y., et al. Intrinsic retroviral reactivation in human preimplantation embryos and pluripotent cells. Nature 2015; 522:221–225

45. Li, M., Radvanyi, L., Yin, B., Rycaj, K., Li, J., Chivukula, R., Lin, K., Lu, Y., Shen, J., Chang, D.Z., et al. Downregulation of Human Endogenous Retrovirus Type K (HERV-K) Viral env RNA in Pancreatic Cancer Cells Decreases Cell Proliferation and Tumor Growth. Clinical cancer research: an official journal of the American Association for Cancer Research 2017; 23:5892–5911

46. Roulois, D., Loo Yau, H., Singhania, R., Wang, Y., Danesh, A., Shen, S.Y., Han, H., Liang, G., Jones, P.A., Pugh, T.J., et al. DNA-Demethylating Agents Target Colorectal Cancer Cells by Inducing Viral Mimicry by Endogenous Transcripts. Cell 2015; 162:961–973

47. Chiappinelli, K.B., Strissel, P.L., Desrichard, A., Li, H., Henke, C., Akman, B., Hein, A., Rote, N.S., Cope, L.M., Snyder, A., et al. Inhibiting DNA Methylation Causes an Interferon Response in Cancer via dsRNA Including Endogenous Retroviruses. Cell 2017; 169:361

48. Schoggins, J.W., MacDuff, D.A., Imanaka, N., Gainey, M.D., Shrestha, B., Eitson, J.L., Mar, K.B., Richardson, R.B., Ratushny, A.V., Litvak, V., et al. Pan-viral specificity of IFN-induced genes reveals new roles for cGAS in innate immunity. Nature 2014; 505:691–695

49. Seth, R.B., Sun, L., Ea, C.K., and Chen, Z.J. Identification and characterization of MAVS, a mitochondrial antiviral signaling protein that activates NF-kappaB and IRF 3. Cell 2005; 122:669–682

50. Kong, M., Guo, L., Xu, W., He, C., Jia, X., Zhao, Z., and Gu, Z. Aging-associated accumulation of mitochondrial DNA mutations in tumor origin. Life Medicine 2022; 10.1093/lifemedi/lnac014

51. Takahashi, A., Loo, T.M., Okada, R., Kamachi, F., Watanabe, Y., Wakita, M., Watanabe, S., Kawamoto, S., Miyata, K., Barber, G.N., et al. Downregulation of cytoplasmic DNases is implicated in cytoplasmic DNA accumulation and SASP in senescent cells. Nature communications 2018; 9:1249

52. Watanabe, S., Kawamoto, S., Ohtani, N., and Hara, E. Impact of senescence-associated secretory phenotype and its potential as a therapeutic target for senescence-associated diseases. Cancer science 2017; 108:563–569

53. Tyagi, R., Li, W., Parades, D., Bianchet, M.A., and Nath, A. Inhibition of human endogenous retrovirus-K by antiretroviral drugs. Retrovirology 2017; 14:21

54. Contreras-Galindo, R., Kaplan, M.H., Dube, D., Gonzalez-Hernandez, M.J., Chan, S., Meng, F., Dai, M., Omenn, G.S., Gitlin, S.D., and Markovitz, D.M. Human Endogenous Retrovirus Type K (HERV-K) Particles Package and Transmit HERV-K–Related Sequences. Journal of virology 2015; 89:7187–7201

55. Hindson, B.J., Ness, K.D., Masquelier, D.A., Belgrader, P., Heredia, N.J., Makarewicz, A.J., Bright, I.J., Lucero, M.Y., Hiddessen, A.L., Legler, T.C., et al. High-throughput droplet digital PCR system for absolute quantitation of DNA copy number. Anal Chem 2011; 83:8604–8610

56. Wang, T., Medynets, M., Johnson, K.R., Doucet-O’Hare, T.T., DiSanza, B., Li, W., Xu, Y., Bagnell, A., Tyagi, R., Sampson, K., et al. Regulation of stem cell function and neuronal differentiation by HERV-K via mTOR pathway. Proceedings of the National Academy of Sciences of the United States of America 2020; 117:17842–17853

57. Lee, Y.N., and Bieniasz, P.D. Reconstitution of an infectious human endogenous retrovirus. PLoS pathogens 2007; 3:e10

58. Dewannieux, M., Harper, F., Richaud, A., Letzelter, C., Ribet, D., Pierron, G., and Heidmann, T. Identification of an infectious progenitor for the multiple-copy HERV-K human endogenous retroelements. Genome research 2006; 16:1548–1556

59. Kim, H.S., Takenaka, O., and Crow, T.J. Isolation and phylogeny of endogenous retrovirus sequences belonging to the HERV-W family in primates. The Journal of general virology 1999; 80 (Pt 10):2613–2619

60. Stengel, A., Roos, C., Hunsmann, G., Seifarth, W., Leib-Mosch, C., and Greenwood, A.D. Expression profiles of endogenous retroviruses in Old World monkeys. Journal of virology 2006; 80:4415–4421

61. Mayer, J., Meese, E., and Mueller-Lantzsch, N. Human endogenous retrovirus K homologous sequences and their coding capacity in Old World primates. Journal of virology 1998; 72:1870–1875

62. Wang, F., Zhang, W., Yang, Q., Kang, Y., Fan, Y., Wei, J., Liu, Z., Dai, S., Li, H., Li, Z, et al. Generation of a Hutchinson-Gilford progeria syndrome monkey model by base editing. Protein & cell 2020;11:809–824

63. Zou, Z., Long, X., Zhao, Q., Zheng, Y., Song, M., Ma, S., Jing, Y., Wang, S., He, Y., Esteban, C.R., et al. A Single-Cell Transcriptomic Atlas of Human Skin Aging. Developmental cell 2020; 10.1016/j.devcel.2020.11.002

64. Ruggieri, A., Maldener, E., Sauter, M., Mueller-Lantzsch, N., Meese, E., Fackler, O.T., and Mayer, J. Human endogenous retrovirus HERV-K(HML-2) encodes a stable signal peptide with biological properties distinct from Rec. Retrovirology 2009; 6:17

65. Medstrand, P., and Blomberg, J. Characterization of novel reverse transcriptase encoding human endogenous retroviral sequences similar to type A and type B retroviruses: differential transcription in normal human tissues. Journal of virology 1993; 67:6778–6787

66. Liu, Z., Li, W., Geng, L., Sun, L., Wang, Q., Yu, Y., Yan, P., Liang, C., Ren, J., Song, M., et al. Cross-species metabolomic analysis identifies uridine as a potent regeneration promoting factor. Cell Discovery 2022; 8:6

67. Ren, X., Hu, B., Song, M., Ding, Z., Dang, Y., Liu, Z., Zhang, W., Ji, Q., Ren, R., Ding, J., et al. Maintenance of Nucleolar Homeostasis by CBX4 Alleviates Senescence and Osteoarthritis. Cell reports 2019; 26:3643-3656.e3647

68. Fu, L., Hu, Y., Song, M., Liu, Z., Zhang, W., Yu, F.X., Wu, J., Wang, S., Izpisua Belmonte, J.C., Chan, P., et al. Up-regulation of FOXD1 by YAP alleviates senescence and osteoarthritis. PLoS Biol 2019; 17:e3000201

69. Lei, J., Jiang, X., Li, W., Ren, J., Wang, D., Ji, Z., Wu, Z., Cheng, F., Cai, Y., Yu, Z.R., et al. Exosomes from antler stem cells alleviate mesenchymal stem cell senescence and osteoarthritis. Protein & cell 2021; 10.1007/s13238-021-00860-9

70. Liang, C., Liu, Z., Song, M., Li, W., Wu, Z., Wang, Z., Wang, Q., Wang, S., Yan, K., Sun, L., et al. Stabilization of heterochromatin by CLOCK promotes stem cell rejuvenation and cartilage regeneration. Cell research 2021; 31:187–205

71. Deng, L., Ren, R., Liu, Z., Song, M., Li, J., Wu, Z., Ren, X., Fu, L., Li, W., and Zhang, W. Stabilizing heterochromatin by DGCR8 alleviates senescence and osteoarthritis. Nature communications 2019; 10:1–16

72. Song, M., Belmonte, J.C.I., and Liu, G.H. Age-related cardiopathies gene editing. Aging (Albany NY) 2019; 11:1327–1328

73. De Cecco, M., Criscione, S.W., Peterson, A.L., Neretti, N., Sedivy, J.M., and Kreiling, J.A. Transposable elements become active and mobile in the genomes of aging mammalian somatic tissues. Aging (Albany NY) 2013; 5:867–883

74. De Cecco, M., Criscione, S.W., Peckham, E.J., Hillenmeyer, S., Hamm, E.A., Manivannan, J., Peterson, A.L., Kreiling, J.A., Neretti, N., and Sedivy, J.M. Genomes of replicatively senescent cells undergo global epigenetic changes leading to gene silencing and activation of transposable elements. Aging cell 2013; 12:247–256

75. Patterson, M.N., Scannapieco, A.E., Au, P.H., Dorsey, S., Royer, C.A., and Maxwell, P.H. Preferential retrotransposition in aging yeast mother cells is correlated with increased genome instability. DNA Repair 2015; 34:18–27

76. Li, W., Prazak, L., Chatterjee, N., Gruninger, S., Krug, L., Theodorou, D., and Dubnau, J. Activation of transposable elements during aging and neuronal decline in Drosophila. Nature neuroscience 2013; 16:529–531

77. Douville, R., Liu, J., Rothstein, J., and Nath, A. Identification of active loci of a human endogenous retrovirus in neurons of patients with amyotrophic lateral sclerosis. Annals of neurology 2011; 69:141–151

78. Barbot, W., Dupressoir, A., Lazar, V., and Heidmann, T. Epigenetic regulation of an IAP retrotransposon in the aging mouse: progressive demethylation and de-silencing of the element by its repetitive induction. Nucleic acids research 2002; 30:2365–2373

79. Balestrieri, E., Pica, F., Matteucci, C., Zenobi, R., Sorrentino, R., Argaw-Denboba, A., Cipriani, C., Bucci, I., and Sinibaldi-Vallebona, P. Transcriptional activity of human endogenous retroviruses in human peripheral blood mononuclear cells. BioMed research international 2015; 2015:164529

80. Nexo, B.A., Villesen, P., Nissen, K.K., Lindegaard, H.M., Rossing, P., Petersen, T., Tarnow, L., Hansen, B., Lorenzen, T., Horslev-Petersen, K., et al. Are human endogenous retroviruses triggers of autoimmune diseases? Unveiling associations of three diseases and viral loci. Immunologic research 2016; 64:55–63

81. Freimanis, G., Hooley, P., Ejtehadi, H.D., Ali, H.A., Veitch, A., Rylance, P.B., Alawi, A., Axford, J., Nevill, A., Murray, P.G., et al. A role for human endogenous retrovirus-K (HML-2) in rheumatoid arthritis: investigating mechanisms of pathogenesis. Clinical and experimental immunology 2010; 160:340–347

82. Garcia-Montojo, M., Fathi, S., Norato, G., Smith, B.R., Rowe, D.B., Kiernan, M.C., Vucic, S., Mathers, S., van Eijk, R.P.A., Santamaria, U., et al. Inhibition of HERV-K (HML-2) in amyotrophic lateral sclerosis patients on antiretroviral therapy. Journal of the neurological sciences 2021; 423:117358

83. Gold, J., Rowe, D.B., Kiernan, M.C., Vucic, S., Mathers, S., van Eijk, R.P.A., Nath, A., Garcia Montojo, M., Norato, G., Santamaria, U.A., et al. Safety and tolerability of Triumeq in amyotrophic lateral sclerosis: the Lighthouse trial. Amyotrophic lateral sclerosis & frontotemporal degeneration 2019; 20:595–604

84. Garcia-Perez, J.L., Morell, M., Scheys, J.O., Kulpa, D.A., Morell, S., Carter, C.C., Hammer, G.D., Collins, K.L., O’Shea, K.S., Menendez, P., et al. Epigenetic silencing of engineered L1 retrotransposition events in human embryonic carcinoma cells. Nature 2010; 466:769–773

85. Liu, E.Y., Russ, J., Cali, C.P., Phan, J.M., Amlie-Wolf, A., and Lee, E.B. Loss of Nuclear TDP-43 Is Associated with Decondensation of LINE Retrotransposons. Cell reports 2019; 27:1409–1421 e1406

86. Yegorov, Y.E., Akimov, S.S., Hass, R., Zelenin, A.V., and Prudovsky, I.A. Endogenous beta-galactosidase activity in continuously nonproliferating cells. Experimental cell research 1998; 243:207–211

87. Imai, Y., Takahashi, A., Hanyu, A., Hori, S., Sato, S., Naka, K., Hirao, A., Ohtani, N., and Hara, E. Crosstalk between the Rb pathway and AKT signaling forms a quiescence-senescence switch. Cell reports 2014; 7:194–207

88. Ma, S., Sun, S., Li, J., Fan, Y., Qu, J., Sun, L., Wang, S., Zhang, Y., Yang, S., Liu, Z., et al. Single-cell transcriptomic atlas of primate cardiopulmonary aging. Cell research 2021; 31:415–432

89. Schneider, C.A., Rasband, W.S., and Eliceiri, K.W. NIH Image to ImageJ: 25 years of image analysis. Nat Methods 2012; 9:671–675

90. Li, H., Handsaker, B., Wysoker, A., Fennell, T., Ruan, J., Homer, N., Marth, G., Abecasis, G., and Durbin, R. The Sequence Alignment/Map format and SAMtools. Bioinformatics 2009; 25:2078–2079

91. Subramanian, A., Kuehn, H., Gould, J., Tamayo, P., and Mesirov, J.P. GSEA-P: a desktop application for Gene Set Enrichment Analysis. Bioinformatics 2007; 23:3251–3253

92. Chen, S., Zhou, Y., Chen, Y., and Gu, J. fastp: an ultra-fast all-in-one FASTQ preprocessor. Bioinformatics 2018; 34:i884–i890

93. Kim, D., Langmead, B., and Salzberg, S.L. HISAT: a fast spliced aligner with low memory requirements. Nat Methods 2015; 12:357–360

94. Anders, S., Pyl, P.T., and Huber, W. HTSeq—a Python framework to work with high-throughput sequencing data. Bioinformatics 2015; 31:166–169

95. Liao, Y., Smyth, G.K., and Shi, W. featureCounts: an efficient general purpose program for assigning sequence reads to genomic features. Bioinformatics 2014; 30:923–930

96. Dobin, A., Davis, C.A., Schlesinger, F., Drenkow, J., Zaleski, C., Jha, S., Batut, P., Chaisson, M., and Gingeras, T.R. STAR: ultrafast universal RNA-seq aligner. Bioinformatics 2013; 29:15–21

97. Jin, Y., Tam, O.H., Paniagua, E., and Hammell, M. TEtranscripts: a package for including transposable elements in differential expression analysis of RNA-seq datasets. Bioinformatics 2015; 31:3593–3599

98. Love, M.I., Huber, W., and Anders, S. Moderated estimation of fold change and dispersion for RNA-seq data with DESeq2. Genome Biol 2014; 15:

99. Xi, Y., and Li, W. BSMAP: whole genome bisulfite sequence MAPping program. BMC Bioinformatics 2009; 10:232

100. Thorvaldsdottir, H., Robinson, J.T., and Mesirov, J.P. Integrative Genomics Viewer (IGV): high-performance genomics data visualization and exploration. Brief Bioinform 2013; 14:178–192

101. Zhang, W., Zhang, S., Yan, P., Ren, J., Song, M., Li, J., Lei, J., Pan, H., Wang, S., Ma, X., et al. A single-cell transcriptomic landscape of primate arterial aging. Nature communications 2020; 11:2202

102. Li, J., Zheng, Y., Yan, P., Song, M., Wang, S., Sun, L., Liu, Z., Ma, S., Izpisua Belmonte, J.C., Chan, P., et al. A single-cell transcriptomic atlas of primate pancreatic islet aging. National science review 2021; 8:nwaa127

103. Balaj, L., Lessard, R., Dai, L., Cho, Y.J., Pomeroy, S.L., Breakefield, X.O., and Skog, J. Tumour microvesicles contain retrotransposon elements and amplified oncogene sequences. Nature communications 2011; 2:180

104. Hu, H., Ji, Q., Song, M., Ren, J., Liu, Z., Wang, Z., Liu, X., Yan, K., Hu, J., Jing, Y., et al. ZKSCAN3 counteracts cellular senescence by stabilizing heterochromatin. Nucleic acids research 2020; 48:6001–6018

105. Padmanabhan Nair, V., Liu, H., Ciceri, G., Jungverdorben, J., Frishman, G., Tchieu, J., Cederquist, G.Y., Rothenaigner, I., Schorpp, K., Klepper, L., et al. Activation of HERV-K(HML-2) disrupts cortical patterning and neuronal differentiation by increasing NTRK3. Cell stem cell 2021; 28:1566-1581 e1568

106. Nair, V.P., Mayer, J., and Vincendeau, M. A protocol for CRISPR-mediated activation and repression of human endogenous retroviruses in human pluripotent stem cells. STAR Protoc 2022; 3:101281–101281

107. Dewannieux, M., Blaise, S., and Heidmann, T. Identification of a functional envelope protein from the HERV-K family of human endogenous retroviruses. Journal of virology 2005; 79:15573–15577

108. Segel, M., Lash, B., Song, J., Ladha, A., Liu, C.C., Jin, X., Mekhedov, S.L., Macrae, R.K., Koonin, E.V., and Zhang, F. Mammalian retrovirus-like protein PEG10 packages its own mRNA and can be pseudotyped for mRNA delivery. Science (New York, NY) 2021; 373:882–889

109. Debacq-Chainiaux, F., Erusalimsky, J.D., Campisi, J., and Toussaint, O. Protocols to detect senescence-associated beta-galactosidase (SA-betagal) activity, a biomarker of senescent cells in culture and in vivo. Nature protocols 2009; 4:1798–1806

110. Kaufmann, K., Muino, J.M., Osteras, M., Farinelli, L., Krajewski, P., and Angenent, G.C. Chromatin immunoprecipitation (ChIP) of plant transcription factors followed by sequencing (ChIP-SEQ) or hybridization to whole genome arrays (ChIP-CHIP). Nature protocols 2010; 5:457–472

111. Yan, P., Liu, Z., Song, M., Wu, Z., Xu, W., Li, K., Ji, Q., Wang, S., Liu, X., Yan, K., et al. Genome-wide R-loop Landscapes during Cell Differentiation and Reprogramming. Cell reports 2020; 32:107870

